# Single cell RNAseq provides a molecular and cellular cartography of changes to the human endometrium through the menstrual cycle

**DOI:** 10.1101/350538

**Authors:** Wanxin Wang, Felipe Vilella, Pilar Alama, Inmaculada Moreno, Marco Mignardi, Wenying Pan, Carlos Simon, Stephen R. Quake

## Abstract

In a human menstrual cycle, the endometrium undergoes remodeling, shedding, and regeneration, all of which are driven by substantial gene expression changes in the underlying cellular hierarchy. Despite its importance in human fertility and regenerative biology, mechanistic understanding of this unique type of tissue homeostasis remains rudimentary. We characterized the transcriptomic transformation of human endometrium at single cell resolution, dissecting the multidimensional cellular heterogeneity of this tissue across the entire natural menstrual cycle. We profiled the behavior of 6 endometrial cell types, including a previously uncharacterized ciliated epithelial cell type, during four major phases of endometrial transformation, and found characteristic signatures for each cell type and phase. We discovered that human window of implantation opens with an abrupt and discontinuous transcriptomic activation in the epithelia, accompanied with widespread decidualized feature in the stromal fibroblasts. These data reveal signatures in the luminal and glandular epithelia during epithelial gland reconstruction, and suggest a mechanism for adult gland formation.

## Introduction

The human menstrual cycle – with its monthly remodeling, shedding, and regeneration of the endometrium – is not shared with many other species. Similar cycles have only been consistently observed in human, apes, and old world monkeys (Emera et al., 2012; Martin, 2007) and not in any of the model organisms which undergo sexual reproduction such as mouse, zebrafish, or fly. This cyclic transformation is executed through dynamic changes in states and interactions of multiple cell types, including luminal and glandular epithelial cells, stromal fibroblasts, vascular endothelial cells, and infiltrating immune cells. Although different categorization schemes exist, the transformation can be primarily divided into two major phases by the event of ovulation: the proliferative (pre-ovulatory) and secretory (post-ovulatory) phase (Noyes et al., 1950). During the secretory phase, the endometrium enters a narrow window of receptive state that is both structurally and biochemically ideal for embryo to implant (Croxatto et al., 1978; Wilcox et al., 1999), i.e. the mid-secretory phase or the window of implantation (WOI). To prepare for this state, the tissue undergoes considerable reconstruction in the proliferative phase, during which one of the most essential elements is the formation of epithelial glands (Filant and Spencer, 2014).

Given the broad relevance in human fertility and regenerative biology, a systematic characterization of endometrial transformation across the natural menstrual cycle has been long pursued. Histological characterizations established the morphological definition of menstrual, proliferative, early-, mid-, and late- secretory phases (Noyes et al., 1950). Bulk level transcriptomic profiling advanced the characterization to a molecular and quantitative level (Riesewijk et al., 2003; Ruiz-Alonso et al., 2012) and demonstrated the feasibility of translating the definition into clinical diagnosis of the WOI (Díaz-Gimeno et al., 2011). However, it has been a challenge to derive unbiased or mechanism-linked characterization from bulk readouts due to the uniquely heterogeneous and dynamic nature of endometrium.

The complexity of endometrium is unlike any other tissue: it consists of multiple cell types which vary dramatically in state through a monthly cycle as they enter and exit the cell cycle, remodel, and undergo various forms of differentiation with relatively rapid rates. The notable variance in menstrual cycle lengths within and between individuals (Guo et al., 2006) adds an additional variable to the system. Thus, transcriptomic characterization of endometrial transformation at our current stage of understanding requires that cell types and states be defined with a minimum of bias. High precision characterization and mechanistic understanding of hallmark events, such as the WOI, requires that we study both the static and dynamic aspects of the tissue. Single cell RNAseq provides an ideal platform for these purposes. We therefore performed a systematic transcriptomic delineation of human endometrium across the natural menstrual cycle at single cell resolution.

## Results

To characterize endometrial transformation across the natural human menstrual cycle, we collected endometrial biopsies from 19 healthy and fertile women, 4-27 days after the onset of menstrual bleeding (**Figure S1**). All women were on regular menstrual cycles, with no influence from exogenous hormone or gynecologic pathology. Single cells were captured and cDNA was generated using Fluidigm C1 medium chips. The fraction of reads mapped to the spike-in controls developed by the External RNA Controls Consortium (ERCC) was used as a metric for quality filtering (Method).

### Human endometrium consists of six cell types across the menstrual cycle

Dimensional reduction via t-distributed stochastic neighbor embedding (tSNE) (Maaten and Hinton, 2008) on the top over-dispersed genes (Method) revealed clear segregation of cells into distinct groups (Figure 1A). We defined cell types as segregations that were not time-associated, i.e. groups encompassing cells sampled across the menstrual cycle. Six cell types were thus identified; canonical markers and highly differentially expressed genes enabled straightforward identification of four of these: stromal fibroblast, endothelium, macrophage, and lymphocyte (Figure 1B). The two remaining cell types both express epithelium-associated markers; one of these cell types is characterized by an extensive list of uniquely expressed genes. Functional analysis (Ashburner et al., 2000; Mi et al., 2017; The Gene Ontology Consortium, 2017) revealed that 56% of genes in this list are annotated with a cilium -associated cellular component or biological process (Figure 1C, **Figure S2**), thereby identifying this cell type as “ciliated epithelium”, specifically with motile cilia (Mitchison and Valente, 2017; Zhou and Roy, 2015). We defined the other epithelial cell type as “unciliated epithelium”.

**Figure 1.**
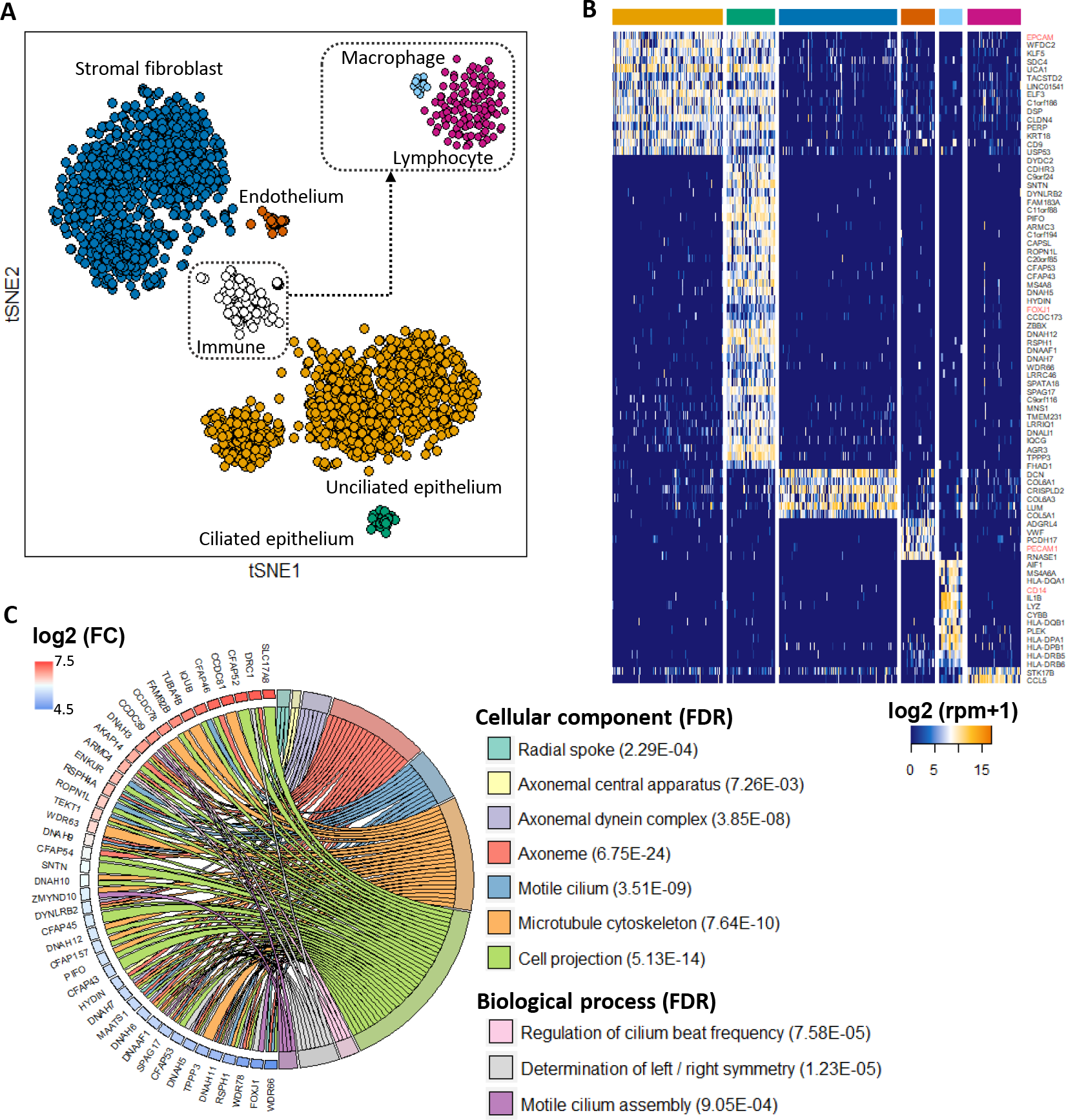
Human endometrium consists of six cell types across the menstrual cycle. (A) Dimension reduction (tSNE) on 2149 single cells from 19 healthy human endometria across the menstrual cycle using top 1000 over-dispersed genes across all cells. Top right inset: tSNE on immune cells using top 1000 over-dispersed genes across immune cells only. Boundaries of cell types were defined by DBSCAN on the 2d-tSNEs. (B) Top discriminatory genes for each identified cell type. Shown are differentially expressed genes (−log_10_(p_adj of a Wilcoxon’s rank sum test)>50, log_2_(FC)>2) that are expressed in >85% cells in the given cell type. For each cell type, genes are ordered, from top to bottom, by the ratio of (% cells within the cell type expressing a gene) and (% cells from other cell types expressing the same gene). In red are canonical markers for the cell type. (C) Cellular components and biological processes enriched in top discriminatory genes for ciliated epithelium. (FC: foldchange, FDR: false discovery rate, p_adj: adjusted p-value) See also Figures S1 and S2

Using RNA and antibody co-staining (Method), we validated previously unannotated discriminatory markers, epithelial lineage identity, and visualized the spatial distribution of ciliated epithelium *in situ*. Four genes were selected for RNA staining: they were identified as highly discriminatory for the cell type (Figure 1B) but either have no previous functional annotation (C11orf88, C20orf85, FAM183A) or are annotated with non-cilia-associated functionality (CDHR3) (**Table S1**). We found consistent co-expression of all four genes with FOXJ1 (canonical master regulator for motile cilia with epithelial lineage identity) antibody staining in both glandular (Figure 2A, C) and luminal (Figure 2B, D) epithelia on day 17 (Figure 2A, B) and day 25 (Figure 2C, D) of the menstrual cycle. The results validated these ciliated cells as an epithelial subpopulation of both luminal and glandular epithelia in healthy human endometrium across the menstrual cycle. This data also demonstrates the consistent discriminatory power of the new markers we identified (Figure 2E) across the cycle. Lastly, the co-expression of these unannotated markers in ciliated cells helps confirm a likely cilia-associated functionality for them and for other unannotated markers we found, which constituted 44% of all markers identified for this cell type (**Figure S2, Table S1**).

**Figure 2.**
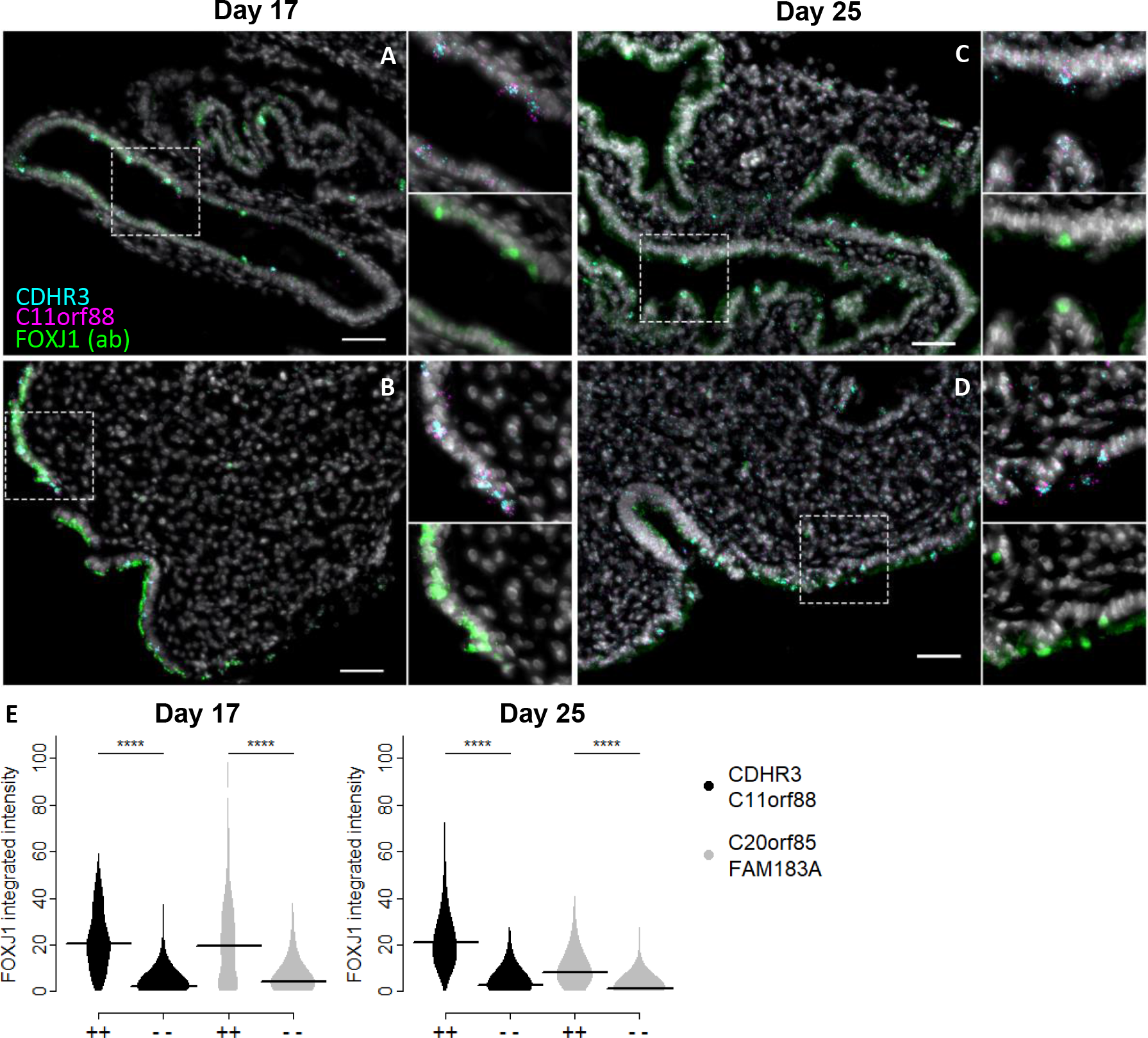
Validation of markers, epithelial lineage, and spatial visualization for endometrial ciliated cells using RNA and antibody co-staining. (A-D) Representative images of human endometrial glands (A, C) and lumen (B, D) on day 17 (A, B) and day 25 (C, D) of the menstrual cycle. (Single CDHR3 and C11orf88 RNA molecules appear as dots in cyan and magenta, respectively. FOXJ1 antibody staining is in green and nuclei in gray. Scale bar: 50 μm. Zoomed-in areas contain triple-expressing cells in the white dashed box in the corresponding panel) (E) Integrated intensity of FOXJ1 antibody for double RNA positive (++) and negative (− −) cells from all images on day 17 (left) and day 25 (right) of the menstrual cycle. (++: cells expressing ≥4 RNA molecules of both markers. Horizonal line: median. ****: p-value of a Wilcoxon’s rank sum test < 0.0001) See also Figures S1 and S2

### Human endometrial transformation consists of four major phases across the menstrual cycle

Samples were taken throughout the menstrual cycle and annotated by the day of menstrual cycle (the number of days after the onset of last menstrual bleeding). While the time variable serves as an informative proxy for assigning endometrial states, it is susceptible to bias due to variances in menstrual cycle lengths between and within women (Guo et al., 2006), and limited in resolution due to variance of cells within an individual. To study transcriptomes of endometrial transformation in an unbiased manner, we performed within-cell type dimension reduction (tSNE) using whole transcriptome data from unciliated epithelia and stromal fibroblasts, respectively. The results revealed four major phases for both cell types, which we refer to as phases 1-4 (**Figure S3A insets**). The four phases were clearly time-associated (**Figure S3A**), confirming the overall validity of the time annotation. Examples where the orders between two women in their phase assignments and time annotation were reversed and cases where cells with the same time annotation were assigned into different phases demonstrate the bias and imitated resolution if we were to use time directly for characterizations (**Figure S3A**).

### Constructing single cell resolution trajectories of menstrual cycle using mutual information based approach

Endometrial transformation over the menstrual cycle is at least in part a continuous process. A model that not only retains phase-wise characteristics but also allows delineation of continuous features between and within phases will enable higher precision characterizations. To build such a model, we used a mutual information (MI) (Tkačik and Walczak, 2011) based approach, such that we exploited the information provided by the time annotation, minimized its limitation noted in the previous section, and accounted for potential continuity between and within phases. Briefly, we enriched for genes that were changing across the menstrual cycle based on the MI between gene expression and time annotation regardless of underlying model of dynamics (Method). In total we obtained 3,198 and 1,156 “time-associated” genes for unciliated epithelia and stromal fibroblasts, respectively (FDR<0.05) (**Figure S3B**). For both cell types, dimensional reduction (tSNE) using time-associated genes revealed the same four major phases that we obtained using unsupervised approach (**Figure S3C, insets**), demonstrating that the MI-based approach reduced the bias of the time annotation to the same extent as unsupervised approach. Meanwhile, the MI-based approach enabled identification of a clear trajectory that connected the phases and was time-associated within phases. We defined the trajectories using the principal curve (Hastie and Stuetzle, 1989) (Figure 3A), and assigned each cell an order along the trajectory based on its projection on the curve (Ji and Ji, 2016; Kim et al., 2016; Marco et al., 2014; Petropoulos et al., 2016), which we refer to as pseudotime (Figure 3A). We observed high correlations between time and pseudotime for both unciliated epithelia and stromal fibroblasts (Figure 3B). The high correlation between pseudotimes of the two cell types from the same woman (Figure 3C) further supports the validity of the trajectories.

**Figure 3.**
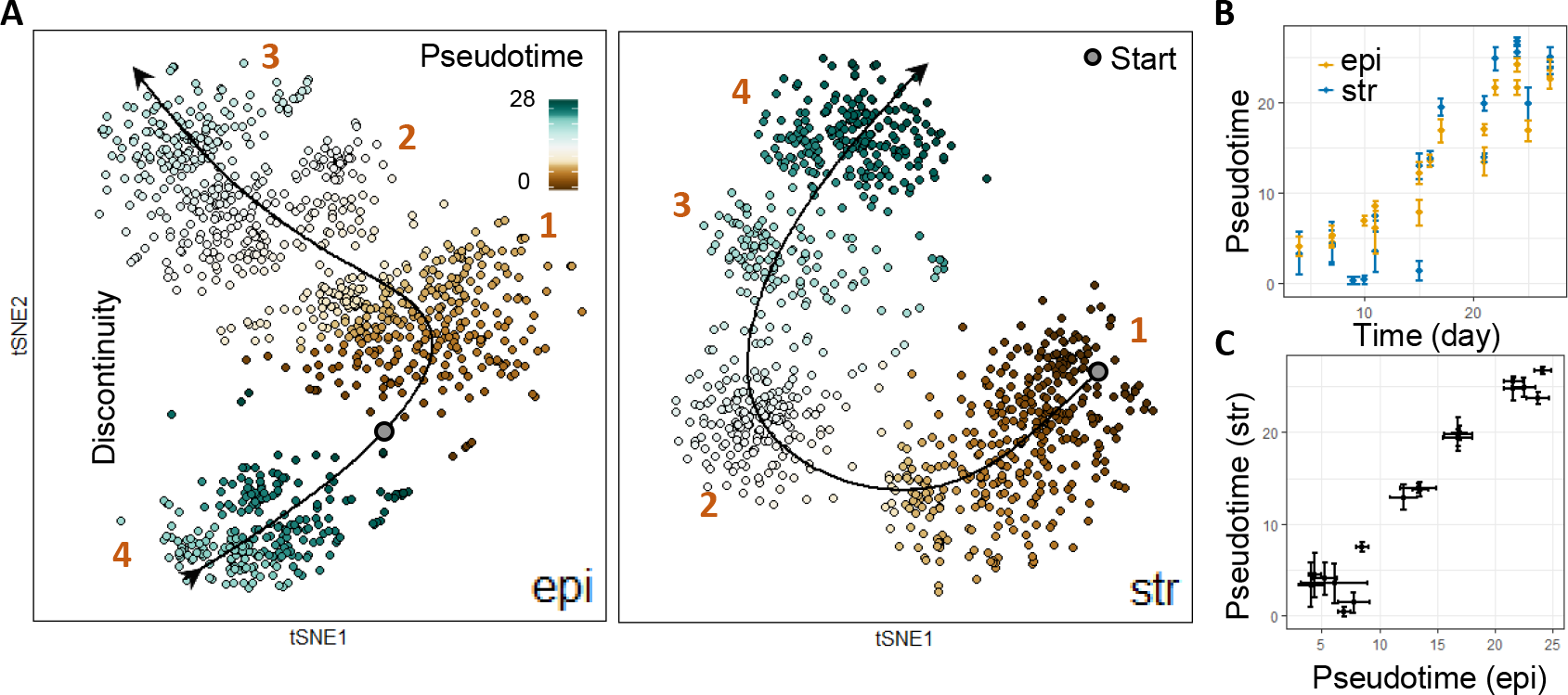
Constructing single cell resolution trajectory of endometrial transformation across the human menstrual cycle. (A) Pseudotime assignment of unciliated epithelia (epi) and stromal fibroblasts (str) across the trajectory of a menstrual cycle. For both cell types, the trajectory was constructed as a principal curve on the 2d-tSNE obtained using “time-associated” genes (see text and Method for details). Pseudotime was assigned as a cell’s order along the trajectory based on its projection on the curve. 1-4: the four major phases consistently identified using either whole transcriptome (Figure S3A) or time-associated genes (Figure S3C). Start: pseudotime=0, assigned based on the clinical definition of the start of a cycle. (B) Correlation of pseudotime and time (day) for epi and str. (C) Correlation of pseudotimes of epi and str from the same woman. (In (B-C), dot and error bar are the median and the median absolute deviation of all epi or str from a woman, respectively. Day: the day of menstrual cycle, i.e. the number of days after the onset of last menstrual bleeding)See also Figures S3-S5

### The WOI opens with an abrupt and discontinuous transcriptomic activation in unciliated epithelium

Interestingly, we observed a notable discontinuity in the trajectory of unciliated epithelia between phase 4 and the preceding phases (Figure 3A, left). This discontinuity was consistently observed regardless of the method we used for dimension reduction (**Figure S4A, S4B**) or feature enrichment (**Figure S4C**). It is unlikely to be an artifact of sampling density given that the involved biopsies were taken with a maximum interval of one day (**Figure S1**) and that a similar discontinuity was not observed in the stromal fibroblast counterpart (Figure 3A, right). To understand the nature of this discontinuity, we explored the genes and their dynamics that contributed to it. Briefly, we identified genes that were dynamically changing along the single-cell trajectories of endometrial transformation by calculating the MI between gene expression and pseudotime, obtaining 1,382 and 527 genes for unciliated epithelia and stromal fibroblasts, respectively (FDR<1E-05, **Figure S5A**). Ordering these genes based on the pseudotime at which their global maximum was estimated to occur (pseudotime_max_, Method) revealed the global features of transcriptomic dynamics across the menstrual cycle (**Figure S5B**). In unciliated epithelia, the dynamics demonstrated an overall continuous feature across phase 1-3, until an abrupt and uniform activation of a gene module marked the entrance into phase 4 (Figure 4A, **S5B left**). Genes in this module included *PAEP*, *GPX3*, and *CXCL14* (Figure 4A), among others which were relatively consistently reported by bulk transcriptomic profilings as overexpressed in the WOI despite notable discrepancies among bulk profiling results(Díaz-Gimeno et al., 2011; Talbi et al., 2006; reviewed by Ruiz-Alonso et al., 2012). Thus, entrance into phase 4 can be identified with the opening of the WOI. Our analysis shows that this transition into the receptive phase of the tissue occurs with an abrupt and discontinuous transcriptomic activation that is uniform among all cells and activated genes in the unciliated epithelia.

### The WOI is characterized by widespread decidualized features in stromal fibroblasts

Unlike their epithelial counterparts, transcriptomic dynamics in stromal fibroblasts demonstrate more stage-wise characteristics, where genes are up-regulated in a modular form, revealing boundaries between phases (Figure 4B, **S5B right**). In phase 4 stromal fibroblasts, the up-regulated gene module includes *DKK1* and *CRYAB*, among a few others that were recapitulated by consensus among bulk analysis and further confirm the identity of WOI (Díaz-Gimeno et al., 2011;Talbi et al., 2006; reviewed by Ruiz-Alonso et al., 2012), although the transition was not as abrupt as in their epithelial counterparts (Figure 4A). In the same module we noticed the decidualization initiating transcriptional factor FOXO1 (Park et al., 2016) and decidualized stromal marker IL15 (Okada et al., 2014). Importantly, while their upregulation in phase 4 was obvious, their expression was already noticeable in phase 3 in a lower percentage of cells and with lower expression level. Decidualization is the transformation of stromal fibroblasts where they change from elongated fibroblast-like cells into enlarged round cells with specific cytoskeleton modifications, playing essential roles for embryo invasion and for pregnancy development (for review see Ramathal. et al., 2010). Our data suggested that this process is initiated before the opening of WOI in a small percentage of stromal fibroblasts, and that at the receptive state of tissue decidualized features are widespread in stromal fibroblasts.

**Figure 4.**
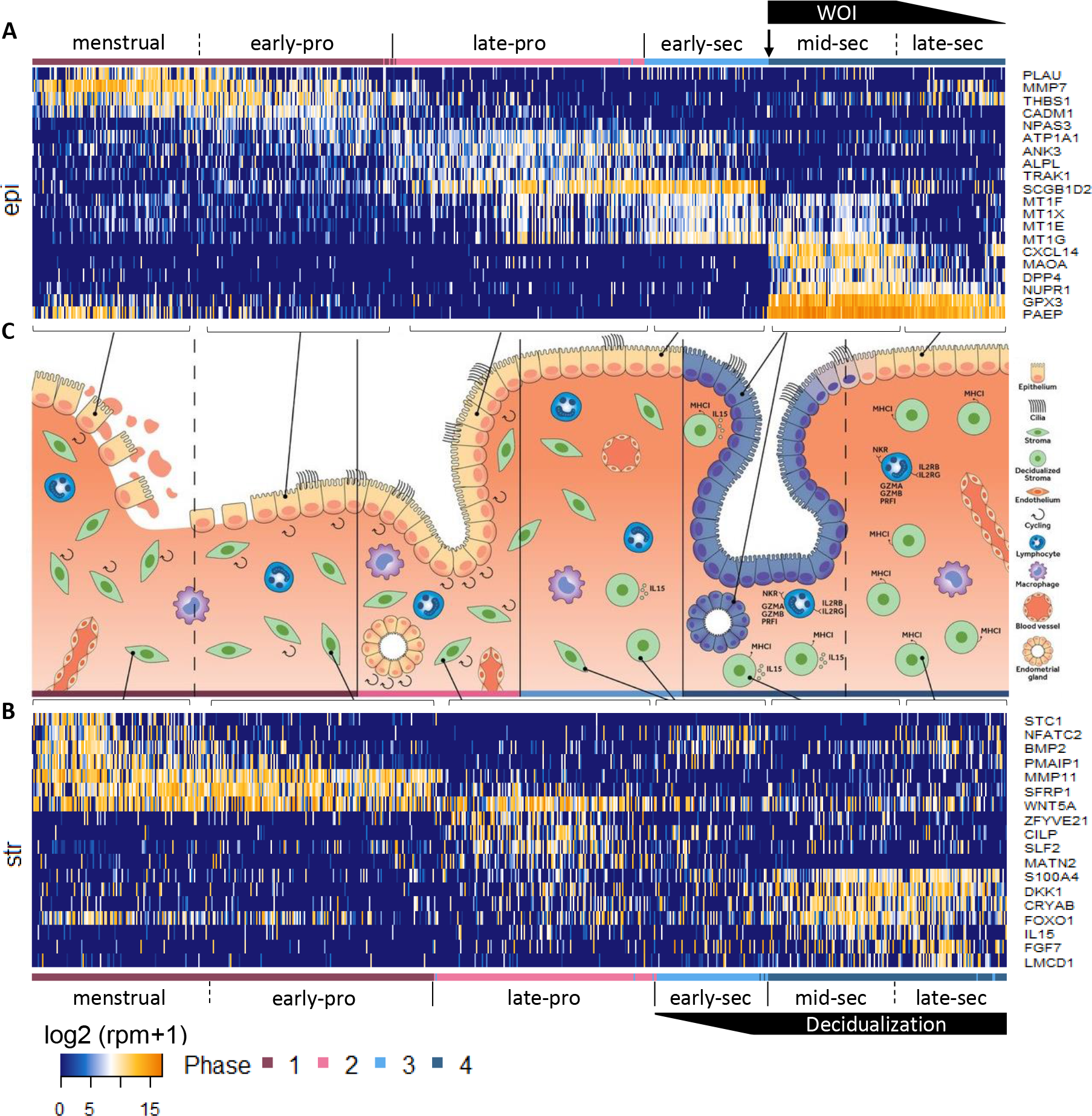
Temporal transcriptome dynamics of endometrial transformation across the human menstrual cycle. Exemplary phase and sub-phase defining genes, and the relationship between transcriptomically defined and histologically defined (canonical) endometrial phases for (A) unciliated epithelia (epi) and (B) stromal fibroblasts (str) across (C) a human menstrual cycle. Shown are genes that differentially expressed (−log_10_(p_adj of a Wilcoxon’s rank sum test)>10, log_2_(FC)>1) in a phase or sub-phase, and are not differentially expressed between luminal and glandular cells in the phase where the gene peaked. Genes were further filtered for their potential to be deconvolutated between unciliated epithelia and stromal fibroblasts in bulk data to obtain those that are temporally in synchrony between the two cell types or those with negligible expression in one cell type across the cycle but significant phase-specific dynamics in another. (Cells (column) were ordered by pseudotime. Dashed line: continuous transition. WOI: window of implantation. pro: proliferative. sec: secretory) See also Figures S6-S9

### The WOI closes with continuous transcriptomic transitions

While the WOI opened up with an abrupt transcriptomic transition in unciliated epithelia, it closed with more continuous transition dynamics (Figure 4A, **S5B left**). Genes expressed in phase 4 unciliated epithelium are featured by three major groups with distinct dynamic characteristics. Group 1 genes (e.g. *PAEP*, *GPX3*) have sustained expression throughout the entire phase 4, and their expression remains noticeable until phase 1 of a new cycle. Group 2 genes (e.g. *CXCL14*, *MAOA*, *DPP4*, and the metallothioneins (*MT1G*, *MT1E*, *MT1F*, *MT1X*)), on the other hand, gradually decrease to zero towards the later part of phase 4, whereas group 3 genes (e.g. *THBS1*, *MMP7*) are upregulated at later part of the phase and their expression is sustained in phase 1 of a new cycle. These characteristics indicate a continuous and gradual transition from mid-secretory to late-secretory phase (Talbi et al., 2006; reviewed by Ruiz-Alonso et al., 2012), and hence the closure of the WOI.

The parallel transition in stromal fibroblasts is also characterized with three similar groups of genes (Figure 4B, **S5B right**) and continuous dynamics. Specifically, we observed a transition towards the later part of phase 4: gradual down-regulation of decidualization–associated genes (e.g. *FOXO1* and *IL15*) and up-regulation of a separate module of genes (e.g*. LMCD1*, *FGF7*). These transitions reveal the final phase of decidualization at the transcriptomic level, which, differing from that during pregnancy, ultimately leads to the shedding of the endometrium in a natural menstrual cycle.

### WOI associated transcriptional regulators are featured with characteristic regulatory roles at the opening and closure of WOI

Cell type identity and cell state are primarily driven by small groups of transcriptional regulators. We therefore sought to identify WOI-associated transcriptional factors (TF) to understand what drives the opening and closure of WOI. We first characterized all TFs that are dynamic across the menstrual cycle (Method) and found for both unciliated epithelia and stromal fibroblasts, these TFs can be primarily assigned to two main categories (**Figure S6A, B, Table S2**), i.e. with 1 or 2 peak(s) of expression detected within one menstrual cycle. Similar to what we observed at whole transcriptome level, the global TF dynamics of the two cell types are notably distinct at the opening of WOI, where in unciliated epithelia a single major discontinuity occurred (**Figure S6A**), whereas in stromal fibroblasts no comparable discontinuity was observed (**Figure S6B**). These, at the level of transcriptional regulators, validated the WOI-associated transcriptomic dynamics described in previous sections.

We next define WOI-associated TFs as those with a peak expression detected after the opening of WOI (**Figure S6C, D**), i.e. the boundary between phase 3 and 4. We further divided these TFs into those 1) that peaked during, and 2) that peaked at the end of phase 4, with the hypothesis that the former are more likely related to the opening of the WOI and the latter the closure. Interestingly, we found that these two groups of TFs are enriched with notably different functional roles. For unciliated epithelia, group 1) TFs are dominated by regulators of early developmental process, especially in differentiation (IRX3, PAX8, MITF, ZBTB20); whereas group 2) TFs include those associated with ER stress (DDIT3) and immediate early genes (FOS, FOSB, JUN). For stromal fibroblasts, group 1) TFs are primarily consisted of regulators of chondrocyte differentiation via cAMP pathways (BHLHE40, ATF3), hence are likely drivers for decidualization, and HIVEP2 - binder to the enhancer of MHC class I genes (discussed more in later sections on immune cells); group 2) TFs include those with roles in ER stress (YBX3, ZBTB16) as well as in regulation of inflammatory (XEBPD) and apoptosis (STAT3). Of note, we highlight the concurrent upregulation of MTF1, which activates the promoter of metallothionein I (**Figure S6C**), with metallothionein I genes (MT1F, 1X, 1E, 1G, Figure 4A) in unciliated epithelia, revealing these heavy metal binding proteins as a key regulatory module associated with WOI.

In summary, our analysis enabled the identification of key drivers for the opening and closure of the human WOI as well as transitions between other major cycle phases (**Figure S6C, D,**top panels). We also highlight the dynamics of nuclear receptors for major classes of steroid hormones (**Figure S6E**), as a special group of TFs mediating the communication between endometrium and other female reproductive organs. Lastly, we performed similar analyses on genes encoding secretory proteins (**Figure S7, Table S3**) and report those associated with the WOI (**Figure S7C, D**).

### The relationship between endometrial phases identified at the transcriptome level is consistent with canonically defined endometrial phases

Since its formalization in 1950 (Noyes et al., 1950), a histological definition of endometrial phases, i.e. the proliferative, early-, mid-, and late-secretory phases, has been used as the gold standard in determining endometrial state. It also usually serves as the ground truth in bulk-based profiling studies in categorizing endometrial phases. Given that there were clear differences between our phase definition and the canonical definition, we investigated the relationship between the two.

Cell mitosis is one of the most distinct features of the pre-ovulatory (proliferative) endometrium, hence the naming of proliferative phase. Thus, to identify the boundary between proliferative and secretory phases, we first explored cell cycle activities across the menstrual cycle. Specifically, we defined endometrial cell cycle associated genes (**Figure S8A**, **B**, Method) and assigned cells into G1/S, G2/M, or non-cycling states (**Figure S8C**, **D**). For both unciliated epithelia and stromal fibroblasts, cell cycling was observed in only a small fraction of cells across the menstrual cycle (**Figure S8C**, **D,** left). This fraction demonstrated phase-associated dynamics, where it was most elevated in phase 1, slightly decreased in phase 2, and almost completely ceased in later phases (**Figure S8C**, **D**, right), indicating that the transition from phase 2 to 3 is between pre-ovulatory to post-ovulatory phases.

To further validate this assignment, we defined characteristic signatures for phase 1-4 (**Table S4,** Method) and identified major hierarchies of biological processes that were enriched by the signatures (**Table S5**, Method). While phase 1 was characterized with processes such as tissue regeneration, e.g. Wnt signaling pathways (unciliated epithelia: epi), tissue morphogenesis (epi), wound healing (stromal fibroblasts: str), and angiogenesis (str) and phase 2 by cell proliferation (epi), phase 3 was dominated by negative regulation of growth (epi) and response to ions (epi) and phase 4 by secretion (epi) and implantation (epi). The transition from a positive to a negative regulation in growth from phase 2 to 3 further confirmed a pre-ovulatory to post-ovulatory transition (Talbi et al., 2006).

Lastly, we used previous bulk tissue analyses to help differentiate the pre-ovulatory and post-ovulatory phases. We reasoned that although bulk data is confounded by the varying proportion of the major cell types, i.e. stromal fibroblasts and unciliated epithelia, bulk and single cell data taken together should have high level of consensus on genes that 1) are in synchrony between the two cell types or 2) have negligible expression in one cell type but significant phase-specific dynamics in another. We therefore identified genes with these characteristics using our single cell data (Figure 4). As expected, among these genes we identified are those that have been consistently reported by bulk studies to be characteristic of canonical endometrial phases, confirming the validity of using them to identify the WOI. Particularly, the upregulation of the metallothioneins (MT1F, X, E, G) from phase 2 and 3 was characteristic of proliferative to early-secretory transition based on bulk reports (Ruiz-Alonso et al., 2012; Talbi et al., 2006). Therefore, considering all of the evidence above, phases 1 and 2 can be identified as pre-ovulatory (proliferative) phases, and phases 3 and 4 as post-ovulatory (secretory) phases. With the anchor provided by the WOI, phase 3 can thus be identified as the early secretory phase.

In phase 1, we observed sub-phases in both unciliated epithelia and stromal fibroblasts that are primarily characterized with genes that are gradually decreasing or increasing towards later part of the phases (Figure 4A, **S5B**). In the unciliated epithelia, the gradually decreasing genes included phase 4 genes (e.g. *PAEP*, *GPX3*), as well as *PLAU*, which activates the degradation of blood plasma proteins. The down-regulation of these genes suggested the end of menstruation, and hence the transition from menstrual to proliferative phase in the canonical definition. Phase 2 can therefore be identified as a second proliferative phase at the transcriptome level. At histological level, transformation in the proliferative endometrium was reported to be featured with morphological changes so gradual that they do not permit the recognition of distinct sub-phases (Noyes et al., 1950). We however have discovered that at the transcriptomic level, proliferative endometrium can be divided into two subphases in both unciliated epithelia and stromal fibroblasts that can be quantitatively identified by transcriptomic signatures (**Figure S9**).

Lastly, we explored interactions between unciliated epithelia and stromal fibroblasts by identifying ligand-receptor pairs that were expressed by the two cell types across the major phases/subphases of the cycle (**Table S6**, Method). We note one major feature within the identified ligand-receptor pairs: they are dominated by a diverse repertoire of extracellular matrix (ECM) proteins paired with integrin receptors, suggesting that ECM-integrin interaction is a major route of communication between the two cell types. We were also able to identify key interactions at the WOI such as between LIF and IL6ST, with LIF being a key gene implicated in endometrial receptivity (Evans et al., 2009, 2016; White et al., 2007).

### Transcriptome signatures in deviating glandular and luminal epithelium supports a mechanism for adult epithelial gland formation

In unciliated epithelia, we noticed further segregation of cells (Figure 5A) in the direction perpendicular to the overall trajectory of the menstrual cycle. To validate this segregation, we independently performed dimension reduction (tSNE) on cells from each of the major phases (**Figure S10A**), excluding genes associated with cell cycles (**Figure S8**). The results confirm the observed segregations when tSNE was done on all unciliated epithelia (Figure 5A).

To identify the nature of this segregation, we performed differential expression analysis and found genes that consistently differentiated the subpopulations across multiple phases (Figure 5B). We examined immunohistochemistry staining of these genes in the Human Protein Atlas (Uhlen et al., 2015) and found that genes upregulated in one population stained intensely in epithelial glands, whereas genes upregulated in the other demonstrated had no to low staining. Moreover, among these genes we found a few that were associated with luminal and glandular epithelia. *ITGA1*, which was reported to be consistently upregulated in glandular epithelia than in luminal epithelia (Lessey et al., 1996), started to differentially express between the two populations at phase 2 and the differential expression persisted for the rest of cycle. *WNT7A*, reported to be exclusively expressed in the luminal epithelium of both humans (Tulac et al., 2003) and mice (Yin and Ma, 2005), overexpressed in the other population in all proliferative phases (Figure 5C); *SVIL*, differentially expressed in the same population in all but phase 4, encodes supervillin, which was associated with microvilli structure responsible for plasma membrane transformation on luminal epithelium (Khurana and George, 2008). Taking the above evidence together, the deviating subpopulations can be identified as the glandular and luminal epithelia.

**Figure 5.**
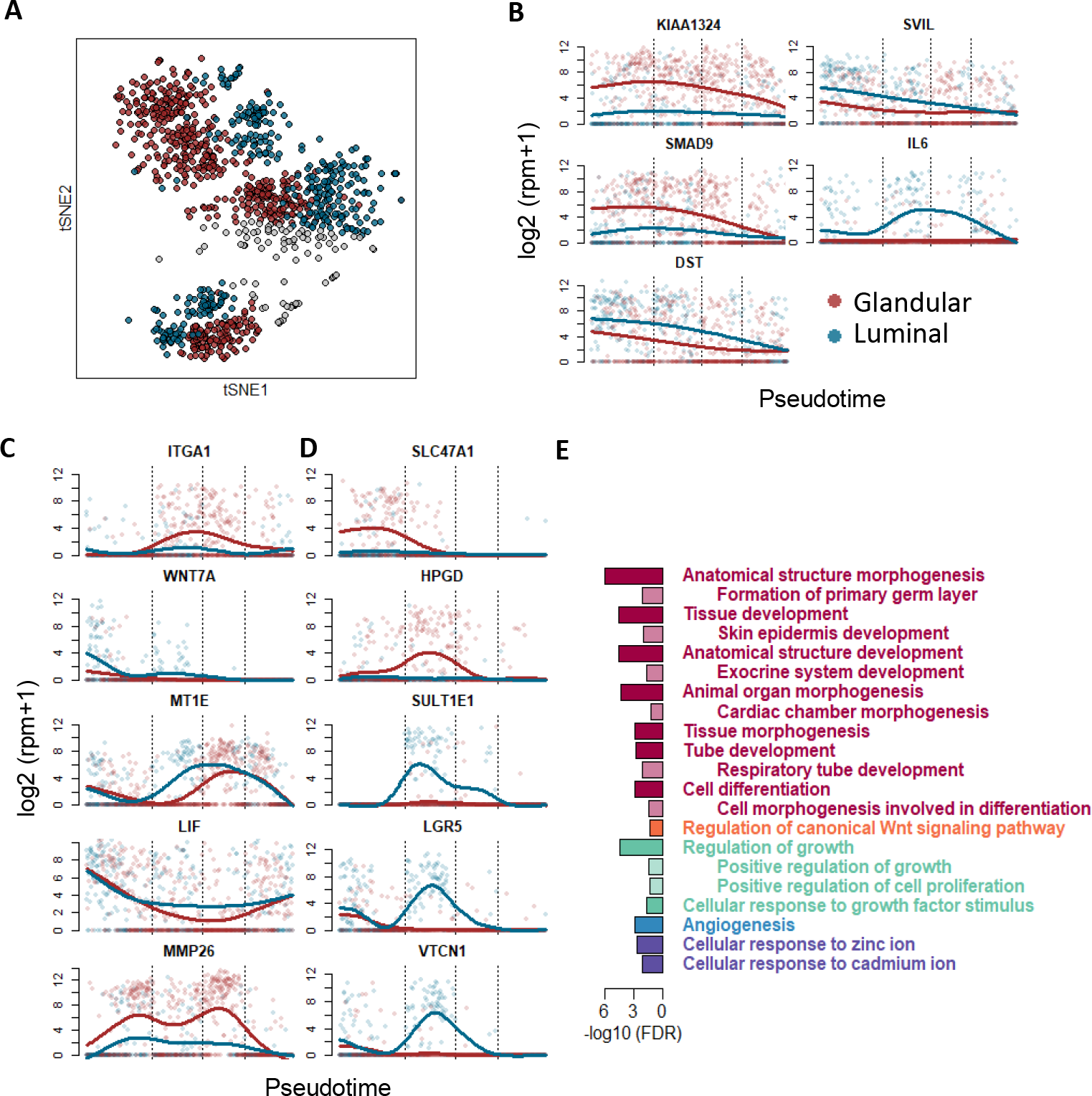
Deviating subpopulations of unciliated epithelia across the human menstrual cycle. (A) Subpopulations of unciliated epithelia. The color-coded classification was based on dimension reduction independently performed for each phase or subphase shown in Figure S10A (gray: cells that are transcriptomically in between the two subpopopulations). (B-D) Dynamics of genes (B) that differentially expressed (−log_10_(p_adj of a Wilcoxon’s rank sum test)>0.05, log_2_(FC)>2) between the two subpopulations across multiple phases, (C) that were previously reported to be implicated in endometrial remodeling or embryo implantation, and (D) that exemplified those that reached maximum differential expression in phase 2. (In (B-D), cells are ordered by pseudotime. Dashed lines: boundaries between the 4 phases) (E) Gene ontology enrichmenched (FDR<0.05) in genes overexpressed in luminal epithelia during proliferative phases. Shown are the enriched GO hierarchies. For each hierarchy, shown are the term with the highest specificity (indented) and the term with the highest significance value (un-indented). See also Figure S10

Within the differentially expressed genes, we also noticed some that were previously reported to be critical for endometrial remodeling and embryo implantation (Figure 5C), and discovered that they were characterized with unique dynamic features. For example, the metallothioneins (*MT1E*, *MT1G*, *MT2A*, *MT1F*) were upregulated in the luminal and glandular cells with a consistent lag in one phase. *LIF*, which was implicated in endometrial receptivity (Evans et al., 2009, 2016; White et al., 2007), was down-regulated in glandular epithelium throughout phase 2, 3, and early phase 4. *MMP26*, a metalloproteinase reported to be up-regulated in proliferative endometrium (Ruiz-Alonso et al., 2012), was differentially expressed in glandular epithelia until phase 4. Of note, we observed no such differential expression in phase-defining genes presented in the earlier sections or housekeeping genes (**Figure S10B**).

Compared to the consistent distinction between the ciliated and unciliated epithelia, the deviation between luminal and glandular epithelia at transcriptome level was subtler and more dynamic: it became noticeable at late phase 1 and was most pronounced in phase 2 (Figure 5A and **Figure S10A**). This observation is further supported by the dynamics of differentially expressing genes such as *HPGD*, *SULT1E1*, *LGR5*, *VTCN1*, and *ITGA1* (Figure 5C, D), among many others (**Figure S10C**), in that the maximum deviation of their expression in luminal and glandular cells was reached in phase 2 (the latest phase before ovulation).

Functional enrichment analysis of genes overexpressed in the luminal epitheliA in proliferative phase revealed extensive enrichments in morphogenesis and tubulogenesis which lead to the development of anatomic structures, as well as morphogenesis at the cellular level which lead to differentiation (Figure 5E). The Wnt signaling pathway, associated with gland formation during the development of the human fetal uterus, was also enriched in this gene group, along with growth, ion transport, and angiogenesis. On the other hand, the most pronounced feature of the glandular subpopulation in the same phase was a consistently higher fraction of cycling cells compared to their luminal counterparts (**Figure S8C**, left). The co-occurrence of the ceasing of cell cycle activity and maximized deviation between the two subpopulations in phase 2 also suggests that the important role proliferation plays in the process.

In addition, we identified a third cell group in the first three biopsies on the pseudotime trajectory (ordered by the median of pseudotime of all cells from a woman) (Figure 5A, **S10A, S12**). This cell group is transcriptomically in between luminal and glandular epithelia (**Figure S10D**), expressing markers from both, suggesting either an intermediate state undergoing transition between two populations or a bipotential progenitor state giving rise to both populations. To explore whether our data supports one state over the other, we examined genes that are overexpressed in this cell group over both luminal and glandular epithelia (**Figure S10E**), where we found genes that are of mesenchymal origin, including *CD90* (*THY1*) and fibrillar collagens (*COL1A1, COL3A1*) as well as transcriptional factors that are associated with transitions between mesenchymal and epithelial states, including *TWIST1*, slug *(SNAI2*) (reviewed by Zeisberg and Neilson, 2009), and *WT1* (reviewed by Miller-Hodges and Hohenstein, 2011). The downregulation of these genes from the ambiguous cell group to unciliated epithelia later in the pseudotime trajectory suggested that it is a bipotential mesenchymal progenitor population that develops into luminal and epithelia through mesenchymal to epithelial transition (MET). In fact, we observed the transition between epithelial and mesenchymal states in cells both at the earliest and the latest timepoints on the pseudotime trajectory (Figure 5A), indicating that the transition peaked both immediately before and after menstruation. This characteristic dynamic is further evidenced by the temporal expression of vimentin (*VIM*), a canonical mesenchymal marker, in unciliated epithelia (**Figure S10F**), where its expression is sustained in phase 1 and 2 (menstrual and proliferative phases), repressed in phase 3 and early phase 4 (early- and mid-secretory phases) and rises again in late phase 4 (late-secretory phase). Surprisingly, several previously proposed markers for endometrial cells with clonogenic and mesenchymal characteristics (reviewed by Evans et al., 2016) including *MCAM* (*CD146*) and *PDGFRB* (Schwab and Gargett, 2007) as well as *SUSD2* (Miyazaki et al., 2012) were not significantly upregulated in the ambiguous cell group.

Adult human endometrial gland formation in menstrual cycles have been proposed to originate from the clonogenic epithelial, or mesenchymal progenitors, or both, in the unshed layer of the uterus (basalis) (Nguyen et al., 2017; W.C. et al., 1997). Our data indicates that endometrial re-epithelization is through MET from mesenchymal progenitors, a process that has been demonstrated in transgenic mouse models (Cousins et al., 2014; Huang et al., 2012; Patterson et al., 2013) but had yet to be observed in human. Our data also shows that following re-epitheliation, endometrial gland reconstruction in adult human endometrium is driven by tubulogenesis in luminal epithelium, which involves the formation of either linear or branched tube structures from a simple epithelial sheet (Hogan and Kolodziej, 2002; Iruela-Arispe and Beitel, 2013)- a mechanism that also contributes to gland formation during the development of human fetal uterus (for review, see Cunha et al., 2017; Robboy et al., 2017). This process is also characterized by proliferation activities that are locally concentrated at glandular epithelium.

### Relative abundance of other endometrial cell types demonstrates phase-associated dynamics

Using the phase definition of unciliated epithelia and stromal fibroblasts, we assigned other endometrial cell types from the same woman into their respective phases and quantified their relative abundance across the cycle (**Figure S11A**). We observed an overall increase in ciliated epithelial cells across proliferative phases and a subsequent decrease in secretory phases as well as a notable rise in lymphocyte abundance from late-proliferative to secretory phases. The change in macrophages was in contrary to previous histological reports (Bonatz et al., 1991; Kamat and Isaacson, 1987); factors such as sampling size for a low abundance cell type and sampling bias in choice of spatial locations in microscopic observations of the tissue may have caused the discrepancy and should be taken into account for future studies.

### Decidualization in natural human menstrual cycle is characterized by direct interplay between lymphocytes and stromal fibroblasts

Infiltrating lymphocytes were reported to play essential roles in decidualization during pregnancy, where they were primarily involved in decidual angiogenesis and regulating trophoblastic invasion (Hanna et al., 2006). Their functions in decidualization during the natural human menstrual cycle, however, remain to be defined. The dramatic increase in lymphocytes abundance in the early secretory phase in our data strongly suggests their involvement in decidualization (**Figure S11A**). We therefore characterized their transcriptomic dynamics across the menstrual cycle to explore their roles and their interactions with other endometrial cell types during decidualization.

Compared to their counterparts in non-decidualized endometrium (i.e. secretory (phase 3) and proliferative phases), lymphocytes in decidualized endometrium (phase 4) in natural menstrual cycle have increased expression of markers that are characteristic of uterine NK cells during pregnancy (*CD69*, *ITGA1*, *NCAM1*/*CD56*) (**Figure S11B**). More interestingly, they express a more diverse repertoire of both activating and inhibitory NK receptors (NKR) responsible for recognizing major histocompatibility complex (MHC) class I molecules (Figure 6A). We observed lymphocytes expressing both NK and T cell markers and those expressing only NK markers (**Figure S11B**), and therefore classified them as “*CD3*+” and “*CD3*−” cells based on their expression of markers characteristic of T cells (**Figure S11B**). Particularly, for both “*CD3*+” and “*CD3*−” cells, a noticeable rise in the fraction of cells expressing *CD56*, the canonical NK marker during pregnancy, occurs as early as the tissue transitioned from proliferative to secretory phase (**Figure S11C**), suggesting again that decidualization was initiated before the opening of the WOI.

**Figure 6.**
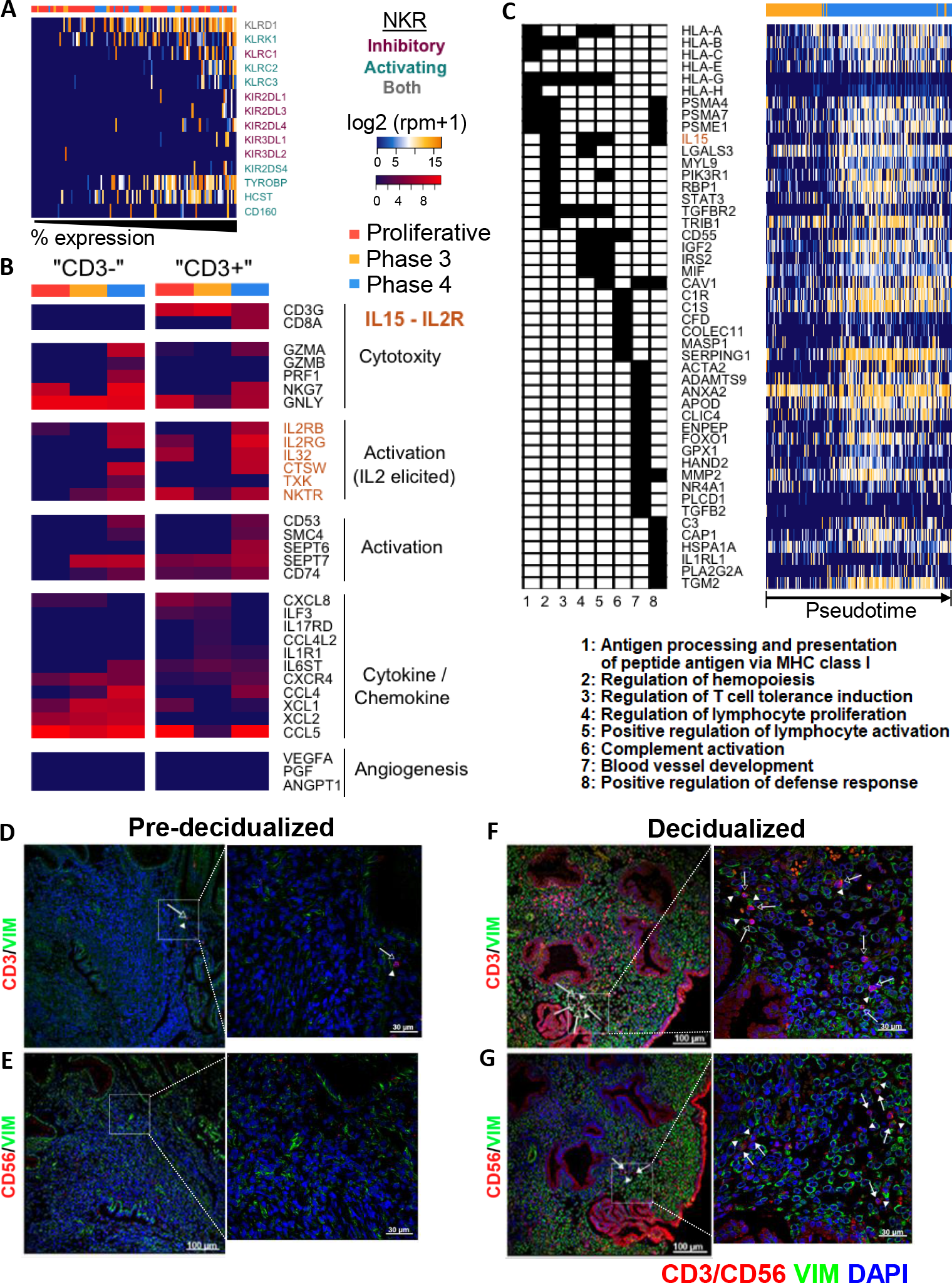
Endometrial lymphocytes across the human menstrual cycle and their interactions with stromal fibroblasts during decidualization. (A) Expression of inhibitory and activating NK receptors (NKR) in endometrial lymphocytes. Cells (columns) were sorted based on percent NKR expressed. (B) Dynamics of genes related to lymphocyte functionality (shown are the medians). “CD3+” and “CD3−” cells are classified based on the expression of markers characteristic of T lymphocytes shown in Figure S11B. (C) Functional annotation (left) and expression (right) of genes that were overexpressed in decidualized stromal fibroblasts (phase 4) that are implicated in immune responses. (D-G) Spatial distribution of CD3 (D, F, open arrow) and CD56 (E, G, arrow) positive immune cells and stromal fibroblasts (arrowhead) before (D, E, day 17) and during (F, G, day 24) decidualization. See also Figure S11

We next identified genes that are dynamically changing in the immune cells across the menstrual cycle and characterized those that are associated with immune functionality (Figure 6B). In “*CD3*−” cells, we observed a significant rise in cytotoxic granule genes in decidualized endometrium (phase 4), with the exception of *GNLY*. In “*CD3*+” cells, this rise in cytotoxic potential is manifested by an increase in *CD8*, while the elevation in cytotoxic granule genes is only moderate. For both “*CD3*+” and “*CD3*−” cells, the increase in IL2 receptors expression is noticeable in phase 4. Equally notable are genes involved in IL2 elicited cell activation. As for the cytokine/chemokine repertoire, “*CD3*−” cells in decidualized endometrium express a high level of chemokines. Their “*CD3*+” counterparts, although expressing a more diverse cytokine repertoire, demonstrate much lower chemokine expression. Lastly, both “*CD3*+” and “*CD3*−” cells in decidualized endometrium have negligible expression in angiogenesis associated genes (Figure 6B), contrary to their counterparts during pregnancy.

Intriguingly, decidualized stromal fibroblasts upregulate immune-related genes that reciprocated those upregulated in phase 4 immune cells. With the diversification of NKR observed in immune cells in the decidualized endometrium (Figure 6A), we observed an overall elevation in MHC class I genes in decidualized stromal fibroblasts (Figure 6C), including *HLA-A* and *HLA-B*, which are recognized by activating NKR, as well as *HLA-G*, recognized by inhibitory NKR. Worth noting was concurrent upregulation of HIVEP2 (**Figure S6D**), a TF responsible for MHC class I genes upregulation. With the IL2-elicited activation observed in immune cells in the decidualized endometrium (Figure 6B), we noticed not only the elevation of *IL15* (plays similar roles as *IL2)* in decidualized stromal fibroblasts, as well as IL15-involved pathways that regulate lymphocyte activation and proliferation (Figure 6C, function annotation #4). Lastly, an angiogenesis associated pathway is elevated in decidualized stromal fibroblasts, complementing the lack of this functionality observed NK cells in the same phase.

Using immunofluorescence, we compared the spatial proximity between the identified immune subsets and stromal fibroblasts before (Figure 6D, E) and during (Figure 6F, G) decidualization. We observed notable increase in the number of both CD3+ (Figure 6D, F) and CD56+ (Figure 6E, G) subsets that are in close proximity with stromal fibroblasts during decidualization compared to pre-decidualization, validating the direct interplays between these immune and stromal subsets during decidualization.

## Discussion

In this work, we studied both static and dynamic characteristics of the human endometrium across the menstrual cycle with single cell resolution. At the transcriptomic level, we used an unbiased approach to identify 6 major endometrial cell types, including a ciliated epithelial cell type, and four major phases of endometrial transformation. For the unciliated epithelia and stromal fibroblasts, we used high-resolution trajectories to track their remodeling through the menstrual cycle with minimum bias. Based on these fundamental units and structures, we identified and characterized the receptive state of the tissue with high precision and studied the dynamic cellular and molecular transformations that lead to the receptive state.

The use of single cell RNAseq to characterize human endometrium is at an early stage. Using endometrial biopsies, a previous study was only limited to the most abundant stromal fibroblasts (late-secretory phase, Krjutskov et al., 2016). Coincident with our work, the feasibility of generating data from other endometrial cell types was also demonstrated by a group using full-thickness uterus (secretory phase, Wu et al., 2018), but cell types were only analyzed at a single time point on a single patient diagnosed with uterine leiomyoma (a gynecological pathology known to cause menstrual abnormalities). Another coincident study modeled decidualization using in vitro cultures of human endometrial stromal fibroblasts and compared the result to the transition of stromal fibroblasts from mid- to late-secretory phase biopsies (Lucas et al., 2018). In our study, biopsies were sampled from 19 healthy women across the entire menstrual cycle. Each of the reported biological phenotypes was supported by multiple biological replicates (i.e. women, **Figure S12**), such that none of the biological results we reported in the study were due to “individual-specific” results, undersampling, or confounded by pathological conditions.

An important result of our work is the molecular characterization of the ciliated epithelium as a transcriptomically distinct endometrial cell type; these cells are consistently present but dynamically changing in abundance across the menstrual cycle (**Figure S11A**). Although the existence of ciliated cells in the human endometrium has been speculated upon based on microscope studies since the 1890’s (Benda, 1894), researchers have been hesitant to include them as an endometrial cell type due to two persisting controversies: 1) whether they exist solely due to pathological conditions (Novak and Rutledge, 1948) and 2) whether they persist across the entire menstrual cycle. The controversies have not been satisfyingly resolved by studies in the 1970’s or recently, due to the confounding gynecological conditions of the examined tissue (Ferenczy et al., 1972; Masterton et al., 1975; Wu et al., 2018) and undersampling (Bartosch et al., 2011). In addition, no standardizable features or signatures were available to identify or isolate this cell type from endometrium. In addition to providing strong evidence that this cell type exists in healthy endometrium throughout the menstrual cycle, we have provided a comprehensive transcriptomic signature along with functional annotations which can serve as molecular anchors for future studies.

In general, ciliary motility facilitates the material transport (e.g. fluid or particles). The notable increase of ciliated epithelia in second proliferative phase (**Figure S11A**) suggests that they may play a role in sperm transport towards fallopian tubes through the uterine cavity. Moreover, their epithelial lineage identity and their consistent presence in glandular epithelia, as shown by our *in situ* results (Figure 2), suggest they may function as the mucociliary transport apparatus, similar to those in the respiratory tract, to transport the secretions and provide proper biochemical milieu. Further elucidation of this role may facilitate more accurate diagnosis of infertility. In addition, we highlight the notably high fraction of genes (~25%) in the derived signature with no functional annotations (**Figure S2, Table S1**). Co-expression of these genes (Figure 1C, 2). with known cilium-associated genes and their exclusive activation in ciliated epithelium provides evidence for their cilium-associated functionality, e.g. in signal sensing and transduction (Bisgrove and Yost, 2006, PNAS Mao et al), whose dysfunction can lead to both organ-specific diseases and multi-system syndromes (Bisgrove and Yost, 2006; Fliegauf et al., 2007). Thus, functional studies that link the roles of these un-annotated genes with cilia functionality will also facilitate the understanding of this organelle.

We identified the opening of the WOI and discovered unique transcriptomic dynamics accompanying both the entrance and the closure of the WOI. It was previously postulated that a continuous dynamics would better describe the entrance of WOI since human embryo implantation doesn’t seem to be controlled by a single hormonal factor as in mice (Hoversland et al., 1982; Paria et al., 1993), while discontinuous characteristics were also speculated based on morphological observation of plasma membrane transformation (Murphy, 2004). Our data suggest that the WOI opens with an abrupt and discontinuous transcriptomic transition in unciliated epithelia, accompanied by a more continuous transition in stromal fibroblasts. The abruptness of the transition also suggests that it should be possible to diagnose the opening of the WOI with high precision in clinical practices of in vitro fertilization and embryo transfer.

It is intriguing that the mid- and late- secretory phases fall into the same major phase at the transcriptomic level, especially since the physiological differences between mid- (high progesterone level, embryo implantation) and late-secretory phase (progesterone withdraw, preparing for tissue desquamation) seem to be as large as that between early- to mid-secretory phase, if not larger. In fact, the characteristic transition at the closure of the WOI is largely contributed by the same group of genes that contributed to the abrupt opening of the WOI, except that while at the opening their upregulation is rapid and uniform across all cells, at the closure the downregulation was executed less uniformly and across a longer period of time. From a dynamic perspective, the difference suggests that the transition between mid-to-late secretory phases, although in magnitude may be similar to that between early-to-mid secretory phases, is slower in rate, perhaps reflective of the rate of progesterone withdrawal. From a molecular perspective, the less uniform downregulation of genes suggests that the closure of the WOI may be mediated through paracrine factors and cell-cell communications.

The abrupt opening of the WOI also allowed us to elucidate the relationship between the WOI and decidualization. As noted earlier, decidualization is the transformation of stromal fibroblasts that is necessary for pregnancy in both human and mouse, and supports developments of implanted embryo. However, contrary to the mouse where decidualization is triggered by implanting embryo(s)(Cha et al., 2012) and thus occurs exclusively during pregnancy, in human, decidualization occurs spontaneously during natural human menstrual cycles independent of the presence of an embryo (Evans et al., 2016). Thus, the relative timing between the WOI and the initiation of decidualization in human is unclear. While histological observation suggests that decidualization starts around mid-secretory phase, our data indicates that decidualization is initiated before the opening of the WOI, and that at the opening of the WOI, decidualized features are widespread in stromal fibroblasts at the transcriptomic level. This lag of morphological signals relative to transcriptomic signals could result from the delay of phenotypic manifestation after transcription either due to inherent delay between transcription and translation or through post-transcriptional modifications.

We identified transcriptomic signatures in the luminal and glandular epithelia during epithelial gland formation. The original definition of luminal and glandular epithelia was established based on the distinct morphology and physical location between the two. Their distinction at the transcriptome level had not been previously established, and we found markers that differentiate the two across multiple phases of the menstrual cycle. Moreover, we discovered signatures that are differentially up-regulated in glandular and luminal epithelium during the formation of epithelial glands. Epithelial glands create a proper biochemical milieu for embryo implantation and subsequent development of pregnancy. In humans, the mechanism for their reconstruction during proliferative phases, however, is unclear. Previous studies through clonogenic assays reported that the cyclic regeneration of both glandular and luminal epithelia was executed by progenitors with stemness characteristics in the unshed layer of the uterus (basalis) (Huang et al., 2012; Nguyen et al., 2017; W.C. et al., 1997). Our analysis suggests a mechanism that involves MET for re-epithelization, followed by tubulogenesis in the luminal epithelium as well as proliferation activities that were locally concentrated at glandular epithelium for reformation of epithelial glands. Our data however cannot rule out the possibility that cells that re-epitheliate the endometrium are the progeny of previously reported candidates with stemness characteristics.

Lastly, we provide evidence for the direct interplay between stromal fibroblasts and lymphocytes during decidualization in menstrual cycle. Our analysis suggests that during decidualization in the cycling endometrium, stromal fibroblasts are directly responsible for the activation of lymphocytes through IL2-eliciated pathways. The diversification of activating and inhibitory NKR in immune cells and the overall up-regulation of MHC class I molecules in stromal fibroblasts is particularly interesting. During pregnancy, cytotoxic NK cells are tolerant towards the semi-allogeneic fetus (Schmitt et al., 2008). This paradoxical phenomenon was hypothesized to be mediated by 1) the upregulation of non-classical MHC class I molecule (HLA-G) (Apps et al., 2007), the ligand to NK inhibitory receptor, and 2) the downregulation of classical MHC class I molecules (HLA-A, HLA-B) (Moffett-King, 2002; Sivori et al., 2000) that engage with NK activating receptors. Our results demonstrate that similar suppression in NK cells with high cytotoxic potential occurs during natural menstrual cycle, however exerted by decidualized stromal fibroblasts.

## Conclusion

In summary, we systematically characterized the human endometrium across the menstrual cycle from both a static and a dynamic perspective. The high resolution of the data and our analytical framework allowed us to answer previously unresolved questions that are centered on the tissue’s receptivity to embryo implantation. We envision that our findings and the molecular signatures we discovered will provide conceptual foundations and practical molecular anchors for future studies.

## Acknowledgments

The authors would like to thank Hongxu Ding for valuable discussion and advice, Norma Neff and Jennifer Okamoto for sequencing expertise, Feiqiao Brian Yu, Spyros Darmanis, and Fabio Zanini for technical assistance and discussion, and Stanford Cell Science Imaging Facility for assistance in imaging. This study was jointly supported by the March of Dimes, the Chan Zuckerburg Biohub, and MINECO/FEDER SAF-2015-67164-R (CS) (Spanish Government). WW was supported by the Stanford Bio-X Graduate Bowes Fellowship. FV was supported by Miguel Servet Program Type II of ISCIII [CPII18/00020] and FIS project [PI18/00957].

## Author Contributions

WW, WP, FV, CS, and SRQ conceived and designed the study. WW, FV, and IM performed experiments. WW performed single cell experiments, RNAscope experiments and imaging. FV optimized the tissue dissociation protocol. IM performed tissue dissociation and immunofluorescence experiments. PA collected endometrial biopsies. WW and SRQ analyzed the single cell RNAseq data. MM and WW analyzed RNAscope data. WW, FV, CS, and SRQ interpreted the results. WW, FV, CS, and SRQ wrote the manuscript.

## STAR Methods

### CONTACT FOR REAGENT AND RESOURCE SHARING

Further information and requests for resources and reagents should be directed to and will be fulfilled by the Lead Contact, SRQ.

### SUBJECT DETAILS

All procedures involving human endometrium were conducted in accordance with the Institutional Review Board (IRB) guidelines for Stanford University under the IRB code IRB-35448 and IVI Valencia, Spain under the IRB code 1603-IGX-016-CS. Collection of endometrial biopsies was approved by the IRB code 1603-IGX-016-CS. There were no medical reasons to obtain the endometrial biopsies. Healthy ovum donors were recruited in the context of the research project approved by the IRB. Informed written consent was obtained from each donor in her natural menstrual cycle (no hormone stimulation) before an endometrial biopsy was performed. De-identified human endometrium was obtained from women aged 18-34, with regular menstrual cycle (3-4 days every 28-30 days), BMI ranging 19-29 kg/m^2^ (inclusive), negative serological tests for HIV, HBV, HCV, RPR and normal karyotype. Women with the following conditions were excluded from tissue collection: recent contraception (IUD in past 3 months; hormonal contraceptives in past 2 months), uterine pathology (endometriosis, leiomyoma, or adenomyosis; bacterial, fungal, or viral infection), and polycystic ovary syndrome.

## METHOD DETAILS

### Endometrium tissue dissociation and population enrichment

A two-stage dissociation protocol was used to dissociate endometrium tissue and separate it into stromal fibroblast and epithelium enriched single cell suspensions. Prior to the dissociation, the tissue was rinsed with DMEM (Sigma) on a petri dish to remove blood and mucus. Excess DMEM was removed after the rinsing. The tissue was then minced into pieces as small as possible and dissociated in collagenase A1 (Sigma) overnight at 4 °C in a 50 mL falcon tube at horizontal position. This primary enzymatic step dissociates stromal fibroblasts into single cells while leaving epithelial glands and lumen mostly undigested. The resulting tissue suspension was then briefly homogenized and left un-agitated for 10 mins in a 50 mL Falcon tube at vertical position, during which epithelial glands and lumen sedimented as a pellet and stromal fibroblasts stayed suspended in the supernatant. The supernatant was therefore collected as the stromal fibroblast-enriched suspension. The pellet was washed twice in 50 mL DMEM to further remove residual stromal fibroblasts. The washed pellet was then dissociated in 400 μL TrypLE Select (Life technology) for 20 mins at 37°C, during which homogenization was performed via intermittent pipetting. DNaseI (100 μL) was then added to the solution to digest extracellular genomic DNA. The digestion was quenched with 1.5 mL DMEM after 5 min incubation. The resulting cell suspension was then pipetted, filtered through a 50 μm cell strainer, and centrifuged at 1000 rpm for 5 min. The pellet was re-suspended as the epithelium-enriched suspension.

### Single cell capture, imaging, and cDNA generation

For cell suspension of both portions, live cells were enriched via MACS dead cell removal kit (Miltenyi Biotec) following the manufacture’s protocol. The resulting cell suspension was diluted in DMEM into a final concentration of 300-400 cells/μL before being loaded onto a medium C1 chip for mRNA Seq (Fluidigm). Live dead cell stain (Life Technology) was added directly into the cell suspension. Single cell capture, mRNA reverse-transcription, and cDNA amplification were performed on the Fluidigm C1 system using default scripts for mRNA Seq. All capture site images were recorded using an in-house built microscopic system at 20x magnification through phase, GFP, and Y3 channels. 1μL pre-diluted ERCC (Ambion) was added into the lysis mix, resulting in a final dilution factor of 1:80,000 in the mix.

### Single cell RNAseq library generation

Single-cell cDNA concentration and size distribution were analyzed on a capillary electrophoresis-based automated fragment analyzer (Advanced Analytical). Tagmented and barcoded cDNA libraries were prepared only for cells imaged as singlet or empty at the capture site and with > 0.06 ng/uL cDNA generated. Library preparation was performed using Nextera XT DNA Sample Preparation kit (Illumina) on a Mosquito HTS liquid handler (TTP Labtech) following Fluidigm’s single cell library preparation protocol with a 4x scale-down of all reagents. Dual-indexed single-cell libraries were pooled and sequenced in pair-end reads on Nextseq (Illumina) to a depth of 1-2 ×10^6^ reads per cell. bcl2fastq v2.17.1.14 was used to separate out the data for each single cell by using unique barcode combinations from the Nextera XT preparation and to generate *.fastq files.

### Tissue preparation for *in situ* hybridizations

Endometrial tissues were fixed for 24-48 h in 4% paraformaldehyde (PFA) at room temperature, trimmed, embedded in paraffin, and sectioned into 3 µm in thickness onto APES-coated slides.

### Immunofluorescence

Tissue sections were baked at 60 °C for 1h, deparaffined with Histoclear and rehydrated with ethanol series. Antigen retrieval was performed by boiling tissue sections in 10 mM sodium citrate buffer (pH 6.0) for 20 min, followed by immediate cool down in cold water for 10 min. Tissue permeabilization was done with 0.25% Triton X 100 in PBS for 5 min, followed by wash in 0.05% Triton X100 in PBS for 5 min twice. Non-specific binding was blocked with 5% BSA-0.05% Triton X100-4% goat serum in PBS for 1h at room temperature. Tissue sections were then incubated with primary antibodies over night at 4 °C and secondary antibodies for 1h at room temperature. Primary antibodies used and dilution ratios are Vimentin (2 µg/mL, ab8978, Abcam), Prolactin (1:10, PA5-26006, Thermo Fischer Scientific), CD3 (1:100, ab5690, Abcam), CD56 (1:50, ab133345, Abcam). Secondary antibodies used and dilution ratios are: Goat anti-mouse IgG (H+L) Superclonal™ Alexa Fluor 488 (1:200, A27034, Thermo Fischer Scientific) and Goat anti-rabbit IgG (H+L) Superclonal™ Alexa Fluor 555 (1:200, A27039, Thermo Fisher Scientific). All sections were counterstained with 4’, 6’-diamidino-2-phenylindole (DAPI) (Thermo Fisher Scientific) and mounted with Aquatex® (Merck-Millipore). Images were captured with a confocal microscope (FV1000, Olympus) at 20X and 60X magnification with oil immersion and analyzed using Imaris (Bitplane).

### RNAscope for ciliated cells

Combined RNA and antibody *in situ* hybridizations were performed according to the manufacturer’s technical note “RNAscope Multiplex Fluorescent v2 Assay combined with Immunofluorescence” for FFPE samples (Advanced Cell Diagnostics). 15 min and 30 min incubation were used for target retrieval and Protease Plus treatment, respectively. RNA probes (Advanced Cell Diagnostics) with the following channel assignment (C), fluorophore, and dilution in TSA buffer were used: CDHR3 (C1, cyanine 3, 1:1500), C11orf88 (C2, cyanine 5, 1:750); C20orf85 (C1, cyanine 3, 1:1500), FAM183A (C2, cyanine 5, 1:1500). Tissue sections were blocked with SuperBlock (PBS) blocking buffer (Fisher Scientific) for 30 min at room temperature, incubated in anti-human FOXJ1 (1:500, eBioscience) over night at 4 °C and goat anti-mouse IgG secondary antibody (1:500, Life Technologies) for 2 h at room temperature. All sections were mounted with Prolong Diamond Antifade Mountant (Thermo Fisher Scientific). Imaging was carried out on an Axio-plan epifluorescence microscope equipped with an Axiocam 506 mono camera (Zeiss) using a 20x/0.8 Plan-Apochromat objective (Zeiss). For each sample, 8-10 fields of view were captured with 10-15 z-stacks.

## QUANTIFICATION AND STATISTICAL ANALYSIS

### Single cell RNAseq data analysis

Raw reads in the *.fastq files were trimmed to 75bp using fastqx, aligned to Ensembl human reference genome GRCh38.87 (dna.primary_assembly) using STAR (Dobin et al., 2013) with default parameters, duplicate-removed using picard MarkDuplicates. Aligned reads were converted to counts using HTSeq (Anders et al., 2015) and Ensembl GTF for GRCh38.87 under the setting -m intersection-strict \-s no. Downstream data analysis was performed in R and Java. For each cell, counts were normalized to log transformed reads per million (log2(rpm+1)) by the equation 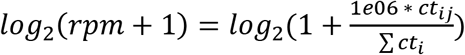 where *i* is for cell *i* and *j* for gene *j*.

### Quality filtering of single cells

For quality filtering, fraction of reads mapped to ERCC (*f*_ERCC_) was used as the quality metric and empirical cumulative distribution of *f*_ERCC_ in empty capture sites recorded on the C1 chip was calculated and used as the null model (*ecdf*_*null*_). Single cells retained for downstream analysis were those with (*ecdf*_*null*_(*f*_*ERCC*_)) < 0.05. 2149 cells were retained for downstream analysis.

### Cell heterogeneity analysis

Over-dispersion of genes was calculated as 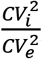, where is 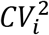 the squared variation of coefficient of gene *i* across cells of interest and 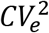 is the expected squared variation of coefficient given mean, fitted using non-ERCC counts. All pairwise distances between cells were calculated as (1-Pearson’s correlation). Dimensional reduction was performed using the R implementation of tSNE (Rtsne).

### Differential expression analysis

To obtain differentially expressed genes for a cell type or state, for each gene, Wilcoxon’s rank sum test (Mann and Whitney, 1947) was performed and fold change (FC, dummy variable = 1E-02) was calculated between cells within a cell type / state and the cells from other cell types / states. P-values obtained from the Wilcoxon’s rank sum test were adjusted for multiple comparisons by Benjamini-Hochberg’s procedure (Benjamini and Hochberg, 1995) to obtain p.adj. To evaluate the “sensitivity” and “specificity” of a gene in identifying a cell type / state, we also calculated the percent of cells within the cell type/state of interest that are expressing the gene (pct_in_) and the percent of cells from other cell types / states expressing the gene (pct_out_), as well as the ratio between the pct_in_ and pct_out_.

### Gene ontology functional enrichment

Functional enrichment analysis was performed using Gene Ontology Enrichment Analysis (http://www.geneontology.org) and each enriched ontology hierarchy (FDR<0.05) was reported with two terms in the hierarchy: 1) the term with the highest significance value and 2) the term with the highest specificity.

### Enrichment of “time-associated” genes via mutual information (MI) based approach

The “time-associatedness” of a gene was calculated as the MI between the expression of a gene and time (or pseudotime) using the Java implementation of ARACNe-AP (Lachmann et al., 2016). For each gene, MI_i_=MI((e_1i_, e_2i_,…, e_ni_),(t_1_, t_2_,…, t_n_)), where i is for gene i, eni is for expression of gene i in cell n, and tn is the time (or pseudotime) annotation of cell n. The statistical significance of the MIi was evaluated using the null model where the time (or pseudotime) annotation was permutated for 1000 times with respect to cells, based on which an empirical cumulative distribution function (ecdf_null,i_) between the expression of gene i and the permutated time (or pseudotime) was constructed using R function ecdf. The p-value for MIi was calculated as (1-ecdf_null,i_(MI_i_)). The p-values were then adjusted for multiple comparisons by Benjamini-Hochberg’s procedure (Benjamini and Hochberg, 1995) to obtain FDR for each gene.

### Smoothing of “time-associated” genes and assignment into characteristic phases

To estimate the pseudotime at which a gene reached maximum expression (pseudotime_max_), smoothing of gene expression was performed with respect to pseudotime using the R function smooth.spline (spar=1) and the pseudotime(s) at which a smoothed curve reached local maximum was estimated using the R function peaks and inflection point estimated using a custom R script.

Characteristic signatures for phase 1-4 (**Table S4**) were identified by assigning each pseudotime-associated gene we identified (**Figure S5A, B**) to the phase where its peak expression occurred (i.e. pseudotime_max_)

### Characterization of dynamics of transcriptional factors (TF) and genes encoding secretory proteins (sec genes) across the menstrual cycle

We define a dynamic TF/sec gene (**Figure S6**) as a “time-associated” gene (**Figure S5B**) that is annotated as a transcriptional regulator/encoding a secretory protein by the Human Protein Atlas (Uhlen et al., 2015). Dynamic TFs/sec genes were first categorized into major groups using hierarchical clustering on smoothed and [0,1] normalized curves. In each group, TFs/sec genes were ordered by the pseudotime where a peak or a major peak (for curves with two peaks) occurred, and ties were broken by the pseudotime where the inflection point occurred.

### Cell cycle analysis

We took a two-step approach in identifying cycling cells and defining endometrium-specific cell cycle signatures. We first used a published gene set encompassing 43 G1/S and 55 G2/M genes (Tirosh et al., 2016), representing the intersection of four previous gene sets (Kowalczyk et al., 2015; Macosko et al., 2015; Whitfield, 2002), and calculated a G1/S and a G2/M score for all single cells in unciliated epithelia and stromal fibroblasts, respectively, following the scoring scheme in (Tirosh et al., 2016). Briefly, cells with at least 2x average expression of either G1/S or G2/M genes than the average of all cells in the respective cell type was assigned as putative cycling cells. We next performed Wilcoxon’s rank sum test (Mann and Whitney, 1947) between the putative cycling cells and the rest of cells in the cell type to enrich for cell-cycle associated transcriptome signatures that were specific to endometrium (**Figure S8A**). To assign cells into G1/S or G2/M phases, we performed dimension reduction on putative cycling cells using the identified signature, which revealed two major populations enriched with known G1/S or G2/M signatures. We assigned genes as either G1/S or G2/M associated by estimating the population at which peak expression of the gene occurred. We then recalculated the G1/S and G2/M scores for each cell using the signature customized for endometrium and finalized the assignment of G1/S and G2/M cells with at least 2x average G1/S or G2/M expression with respect to all cells in that cell type.

### Identification of putative ligand-receptor interactions between unciliated epithelia and stromal fibroblasts

For each identified phase and subphase, the expression of a known ligand or receptor was evaluated as the percent of unciliated epithelia or stromal fibroblasts expressing the genes to obtain p_(epi, j)_ and p_(str, j)_, where j is for phase j. A ligand or receptor is only considered expressed by a cell type in a phase if p is greater than 25%. The interaction between a ligand-receptor pair is established if when a ligand is expressed in one cell type and its known receptor is expressed in the other. The ligand-receptor pairing information was based on the database provided by (Ramilowski et al., 2015). In **Table S6**, ligand-receptor pairs were sorted, from top to bottom, by the level of interaction, quantified as the total number of interactions normalized by the total number of possible interactions between the two cell types within a phase.

### Analysis of RNAscope images

Z-stacks were projected (maximum intensity projection, MIP) using ImageJ. The resulting MIP images were analyzed using CellProfiler 3.0.0 as follows: 1) Correct background by subtracting the lower quartile of the intensity measured from the whole image. 2) Detect cell nuclei using the DAPI channel and cell boundaries using Voronoi distance (25 pixels) from the nuclei. 3) Enhance RNA signals using a tophat filter (5 pixels) and detect signals by intensity threshold (0.004 and 0.002 for Cy3 and Cy5, respectively). 4) Measure antibody intensity for each detected cell. All images were analyzed in the same way, with no image excluded.

## DATA AND SOFTWARE AVAILABILITY

The datasets generated and analyzed in the study are available in the NCBI Gene Expression Omnibus (GEO) and Sequence Read Archive (SRA) and can be accessed upon request. All custom scripts can be accessed upon request to the Lead Contact.

## Supplemental Figure titles and legends

**Figure S1. Number of single cells sampled across the human menstrual cycle**

(Day: the day of menstrual cycle, i.e. the number of days after the onset of last menstrual bleeding)

(Related to Figure 1)

**Figure S2. Classes of functional annotation and their distribution for uniquely expressed genes in ciliated epithelium**

(Related to Figure 1, 2)

**Figure S3. Constructing single cell resolution trajectories of the human menstrual cycle using mutual information (MI) based approach**

(A) Unbiased definition of four major phases of endometrial transformation across the human menstrual cycle via tSNE on all genes detected for unciliated epithelia (epi) and stromal fibroblasts (str) (Inset: phase assignment using Ward’s hierarchical agglomerative clustering)

(B) MI between gene expression and time (red) or permutated time (black) (Genes are ranked by MI)

(C) tSNE using time-associated genes and trajectories of endometrial transformation defined by principal curves (Inset: phase assignment using Ward’s hierarchical agglomerative clustering) (Related to Figure 3)

**Figure S4. Discontinuity between phase 3 and 4 unciliated epithelia is supported by different analysis methods**

Dimension reduction of unciliated epithelia (epi, left) and stromal fibroblasts (str, right) via

(A) principal component analysis (linear) and

(B) multidimensional scaling (non-linear) using whole transcriptome information

(C) tSNE on top 50 principal components obtained via principal component analysis on whole transcriptome information

(Phase 1-4 assignment followed Figure S3C)

(Related to Figure 3)

**Figure S5. Global temporal transcriptome dynamics across the human menstrual cycle**

(A) MI between expressions of pseudotime-associated genes (FDR<1E-05) and pseudotime (red) or permutated pseudotime (black) for unciliated epithelia (epi) and stromal fibroblasts (str)

(B) Dynamics of pseudotime associated genes across the menstrual cycle

(Related to Figure 4)

**Figure S6. Dynamic transcriptional factors (TF) across the menstrual cycle**

(A, B) All pseudotime associated TFs for unciliated epithelia (epi, A) and stromal fibroblasts (str, B) (genes bracketed by red bars are zoomed in C, D)

(C, D) TFs that are associated with the entrance/exit of WOI (bottom) or phase-defining (top) in epi (C) and str (D)

(E) Expression of TFs that are nuclear hormone receptors for estrogen (ESR1), progesterone (PGR), glucocorticoid (NR3C1), and androgen (AR)

(For heatmap, TFs were ordered first by the pseudotime of the major peak and then by the pseudotime of the peak’s inflection point)

(Related to Figure 3, 4)

**Figure S7. Dynamic genes for secretory proteins (sec genes) across the menstrual cycle**

(A, B) All pseudotime associated sec genes for unciliated epithelia (epi, A) and stromal fibroblasts (str, B) (genes bracketed by blue bars are zoomed in C, D)

(C, D) Sec genes that are associated with the entrance/exit of WOI (bottom) in epi (C) and str (D)

(For heatmap, sec genes were ordered following the same strategy as in Figure S6)

(Related to Figure 4)

**Figure S8. Endometrial cell cycle activities across the menstrual cycle**

(A, B) Endometrial G1/S and G2/M signatures for unciliated epithelia (epi, A) and stromal fibroblasts (str, B)

(C, D) Distribution (left) and factional dynamics (right) of cycling cells across major phases of the menstrual cycle (Related to Figure 4)

Figure S9. Top phase-defining genes for the two proliferative phases for unciliated epithelia (epi) and stromal fibroblasts (str)

(Related to Figure 4)

**Figure S9. Top phase-defining genes for the two proliferative phases for unciliated epithelia (epi) and stromal fibroblasts (str)**

(Related to Figure 4)

**Figure S10. Deviating subpopulations of unciliated epithelia and their transcriptomic signature across the human menstrual cycle**

(A) Dimension reduction (tSNE) on unciliated epithelia in major identified phases/sub-phases across the menstrual cycle

(B) Dynamics of phase-defining and housekeeping genes in unciliated epithelial subpopulations across the menstrual cycle (dashed lines: boundaries between the 4 phases)

(C) Dynamics of differentially expressed genes between the two sub-populations in phase 2

(D) Relationship between the ambiguous cell population with luminal and glandular cells in early phase 1. Shown are differentially expressed genes (-log_10_(p_adj of a Wilcoxon’s rank sum test)>0.05, log_2_(FC)>2) between luminal and glandular epithelia in early phase 1. Cells (column) are ordered by the ratio of (average expression of genes upregulated in the luminal) and (average expression of genes upregulated in the glandular)

(E) Genes over-expressed and under-expressed in the ambiguous cell population relative to in luminal and glandular epithelia in early phase 1. Cells (column) are ordered by the ratio of (average expression of genes under-expressed) and (average expression of genes over-expressed)

(F) Expression of vimentin (VIM) in unciliated epithelia

(Related to Figure 5)

**Figure S11. Changes in other endometrial cell types across the human menstrual cycle**

(A) Normalized abundance of other endometrial cell types demonstrated phase-associated dynamics. Normalization was done against the total number of unciliated epithelia (for ciliated epithelium) or stromal fibroblasts (for lymphocyte, endothelium, macrophage) captured for each biopsy

(B) Expression of major lymphoid lineage markers. Cells (columns) were sorted based on percent NK receptors expressed (as in Figure 6A)

(C) Percent CD56+ cells in all “CD3+” and “CD3−” lymphocytes across major phases of the cycle

(Related to Figure 6)

**Figure S12. Data summary**

For each woman:

(A) Relationship between the day of menstrual cycle and her assignment to one of the four major phases (Figure 3) based on unbiased single cell analysis

(B) Total number of single cells analyzed

(C) Distribution of the six cell types

(D) Distribution of glandular and luminal epithelia (Gray: cells in the ambiguous cell population in Figure 5A)

(Each dot (A, B) or each bar (C, D) represents a woman. From left to right, women were ordered, based on the median pseudotime of her stromal fibroblasts and unciliated epithelia)

**Table S1. Classification of unannotated markers for ciliated epithelial cells**

**Table S2. Dynamic transcriptional factors across the menstrual cycle ordered as in Figure S6**

**Table S3. Dynamic genes encoding secretory genes across the menstrual cycle ordered as in Figure S7**

**Table S4. Characteristic pseudotime-associated genes for each major phase identified Table S5. Hierarchies of biological processes enriched in characteristic pseudotome-**

**associated genes for each major phase**

**Table S6. Ligand receptor pairs between unciliated epithelia and stromal fibroblasts identified for each phase and subphase identified**

