## Supplemental Figures for "Single cell RNAseq provides a molecular and cellular cartography of changes to the human endometrium through the menstrual cycle"

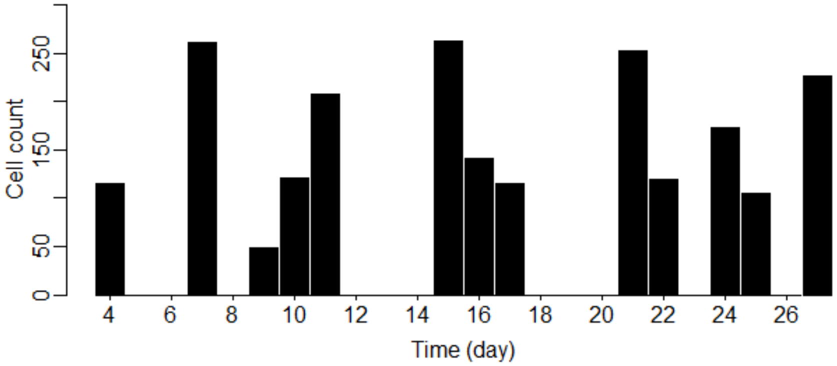


**Figure S1. Number of single cells sampled across the human menstrual cycle**

(Day: the day of menstrual cycle, i.e. the number of days after the onset of last menstrual bleeding)

(Related to Figure 1)

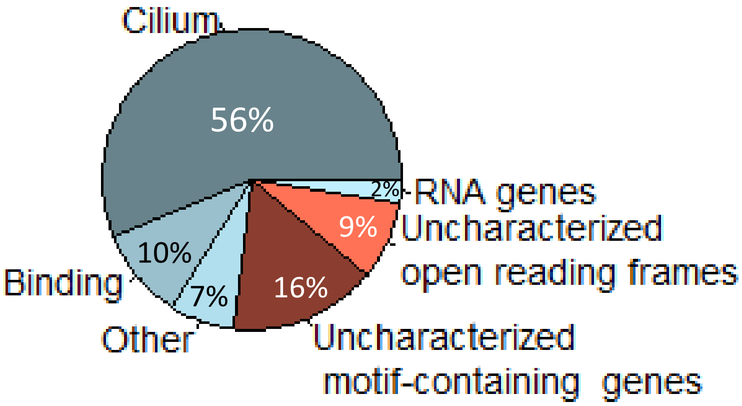


**Figure S2. Classes of functional annotation and their distribution for uniquely expressed genes in ciliated epithelium**

(Related to Figure 1, 2)

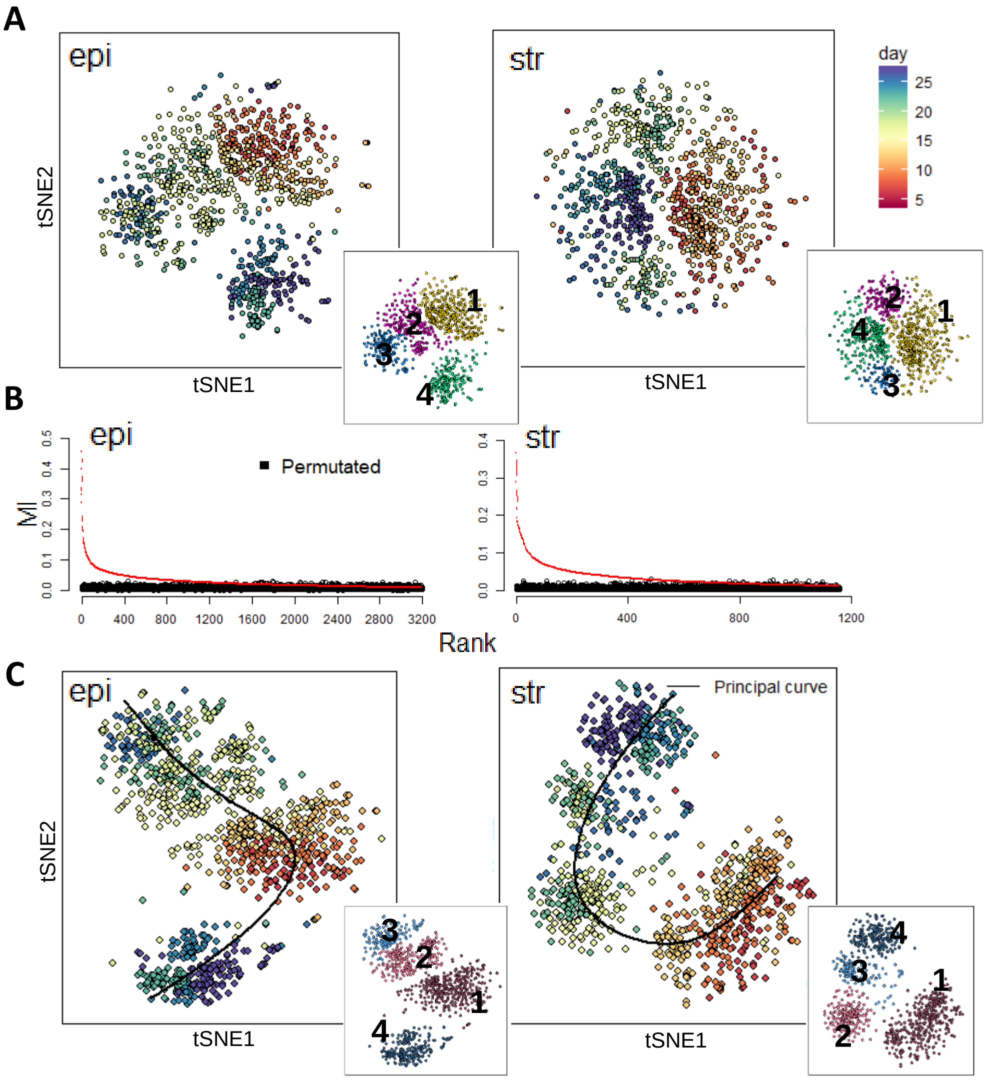


**Figure S3. Constructing single cell resolution trajectories of the human menstrual cycle using mutual information (MI) based approach**

(A) Unbiased definition of four major phases of endometrial transformation across the human menstrual cycle via tSNE on all genes detected for unciliated epithelia (epi) and stromal fibroblasts (str) (Inset: phase assignment using Ward's hierarchical agglomerative clustering)

(B) MI between gene expression and time (red) or permutated time (black) (Genes are ranked by MI)

(C) tSNE using time-associated genes and trajectories of endometrial transformation defined by principal curves (Inset: phase assignment using Ward's hierarchical agglomerative clustering)

(Related to Figure 3)

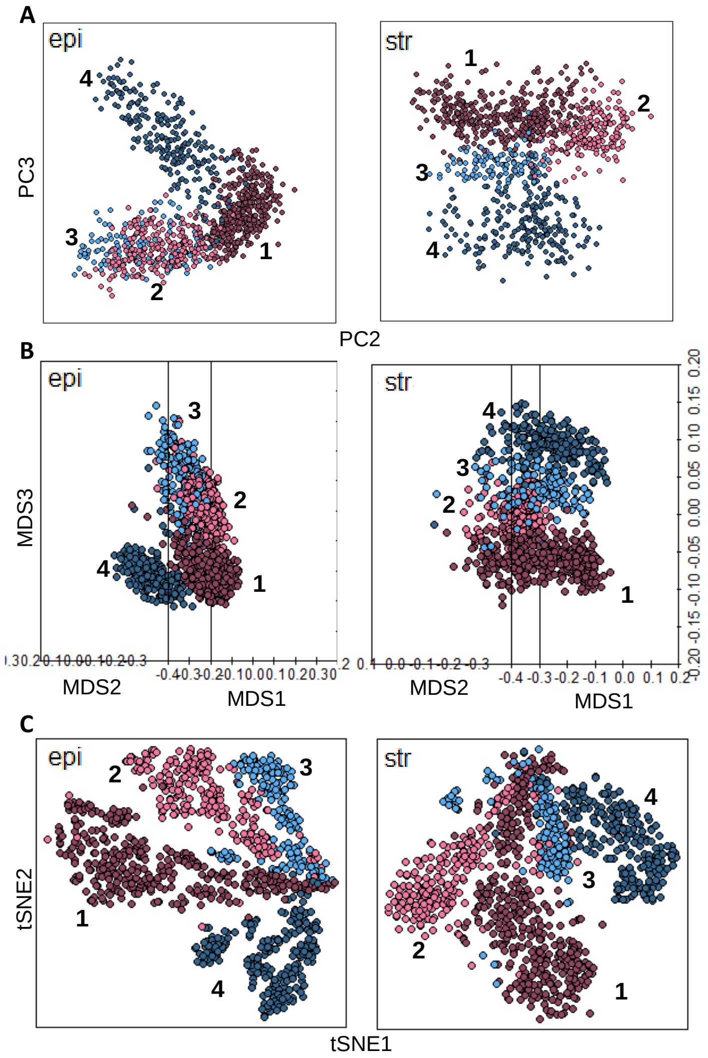


**Figure S4. Discontinuity between phase 3 and 4 unciliated epithelia is supported by different analysis methods**

Dimension reduction of unciliated epithelia (epi, left) and stromal fibroblasts (str, right) via

(A) principal component analysis (linear) and

(B) multidimensional scaling (non-linear) using whole transcriptome information

(C) tSNE on top 50 principal components obtained via principal component analysis on whole transcriptome information

(Phase 1-4 assignment followed Figure S3C)

(Related to Figure 3)

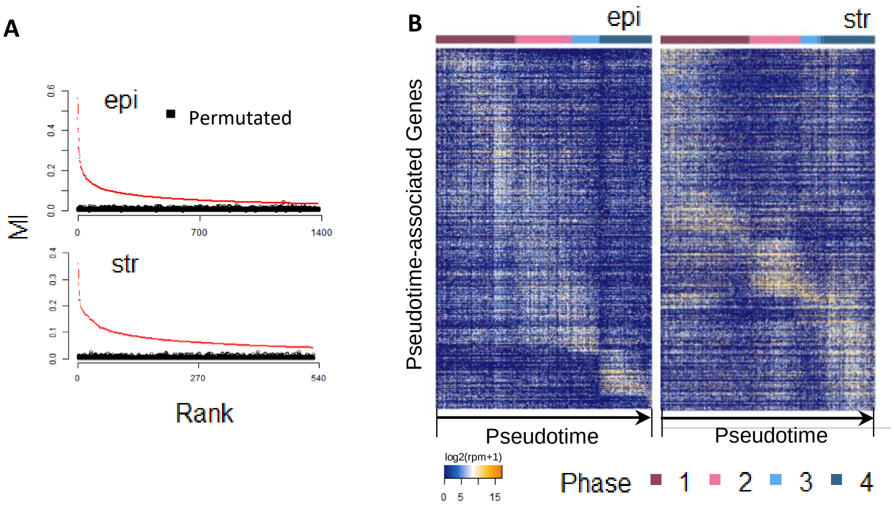


**Figure S5. Global temporal transcriptome dynamics across the human menstrual cycle**

(A) MI between expressions of pseudotime-associated genes (FDR<1E-05) and pseudotime (red) or permutated pseudotime (black) for unciliated epithelia (epi) and stromal fibroblasts (str)

(B) Dynamics of pseudotime associated genes across the menstrual cycle

(Related to Figure 3, 4)

Figure S6

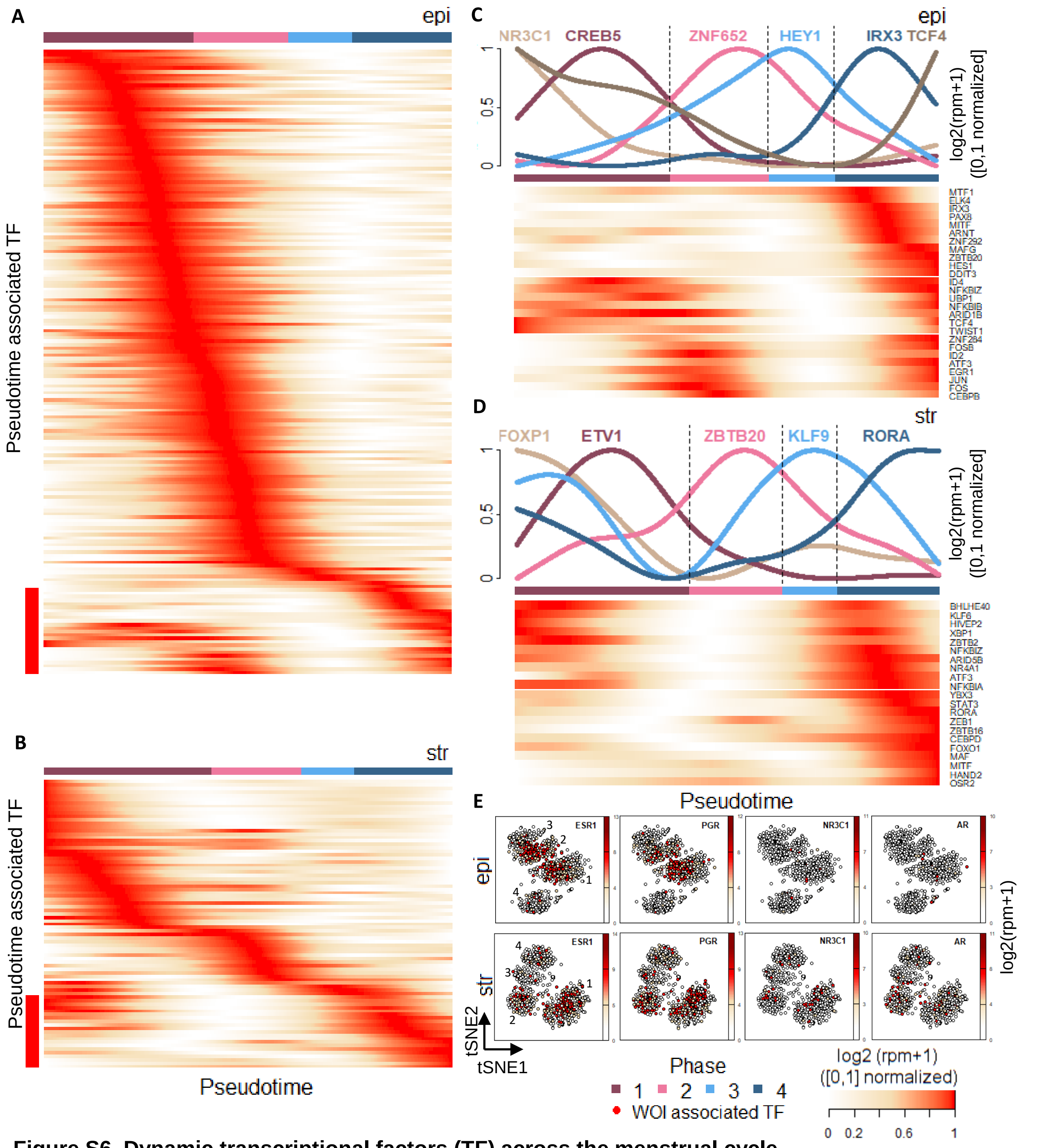


**Figure S6. Dynamic transcriptional factors (TF) across the menstrual cycle**

(A, B) All pseudotime associated TFs for unciliated epithelia (epi, A) and stromal fibroblasts (str, B) (genes bracketed by red bars are zoomed in C, D)

(C, D) TFs that are associated with the entrance/exit of WOI (bottom) or phase-defining (top) in epi (C) and str (D)

(E) Expression of TFs that are nuclear hormone receptors for estrogen (ESR1), progesterone (PGR), glucocorticoid (NR3C1), and androgen (AR)

(For heatmap, TFs were ordered first by the pseudotime of the major peak and then by the pseudotime of the peak's inflection point)

(Related to Figure 4)

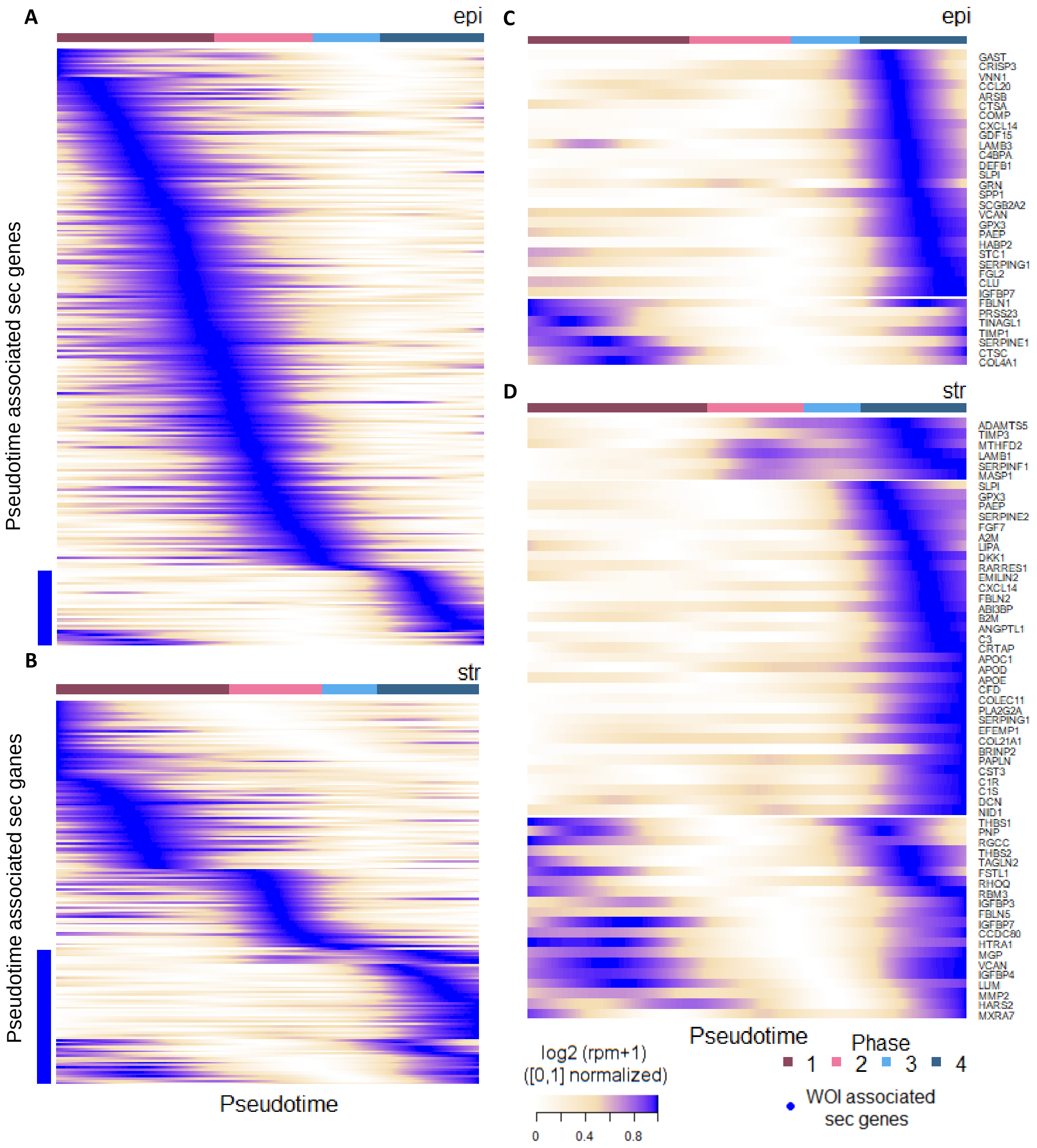


**Figure S7. Dynamic genes for secretory proteins (sec genes) across the menstrual cycle**
(A, B) All pseudotime associated sec genes for unciliated epithelia (epi, A) and stromal fibroblasts (str, B) (genes bracketed by blue bars are zoomed in C, D)
(C, D) Sec genes that are associated with the entrance/exit of WOI (bottom) in epi (C) and str (D)

(For heatmap, sec genes were ordered following the same strategy as in Figure S6)

(Related to Figure 4)

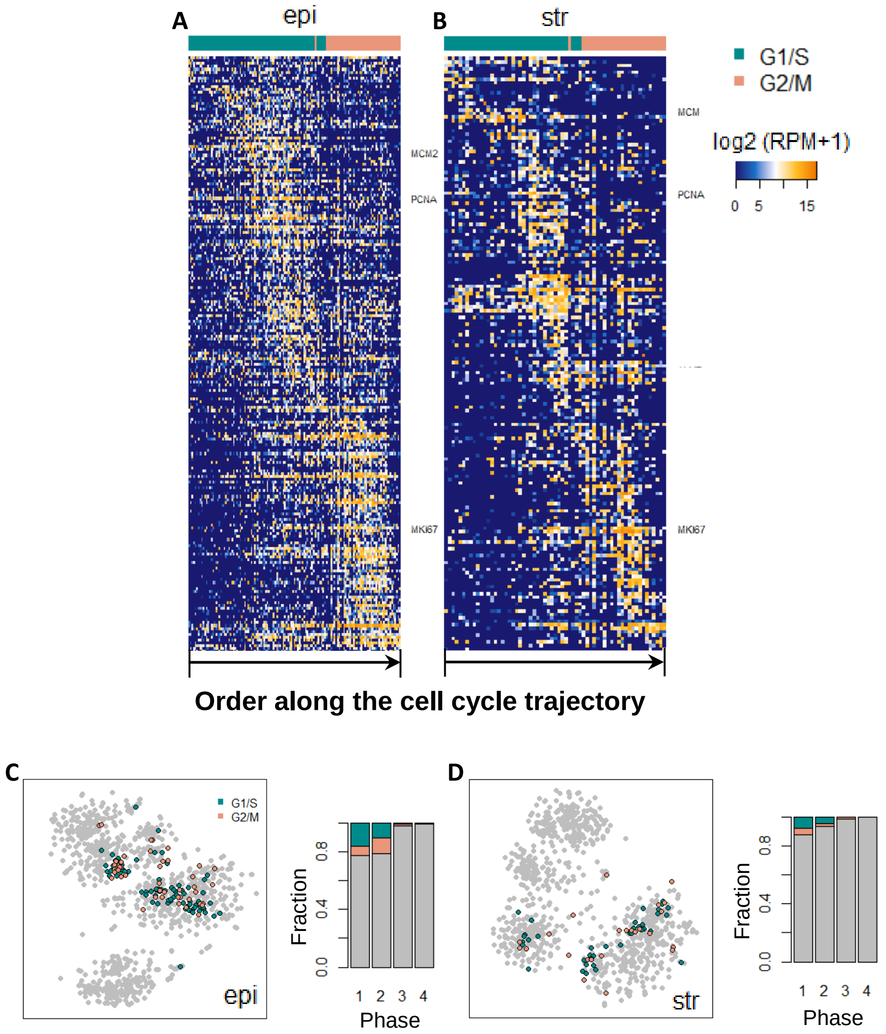


**Figure S8. Endometrial cell cycle activities across the menstrual cycle**
(A, B) Endometrial G1/S and G2/M signatures for unciliated epithelia (epi, A) and stromal fibroblasts (str, B)
(C, D) Distribution (left) and factional dynamics (right) of cycling cells across major phases of the menstrual cycle

(Related to Figure 4)

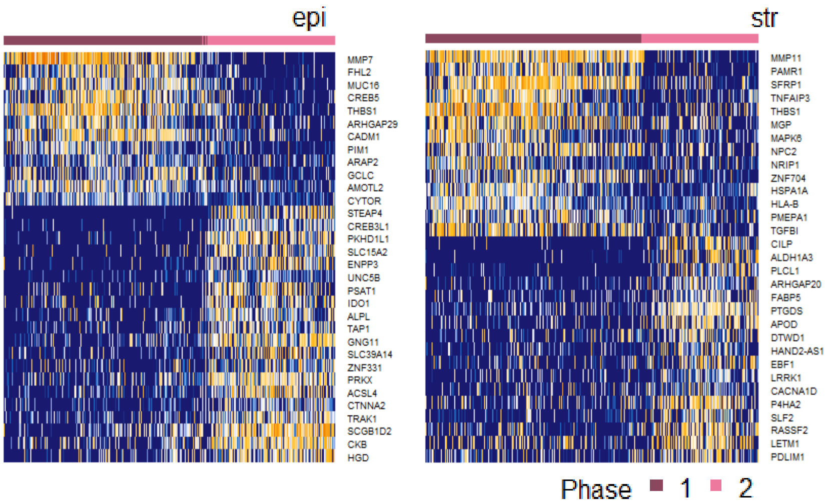


**Figure S9. Top phase-defining genes for the two proliferative phases for unciliated epithelia (epi) and stromal fibroblasts (str)**

(Related to Figure 4)

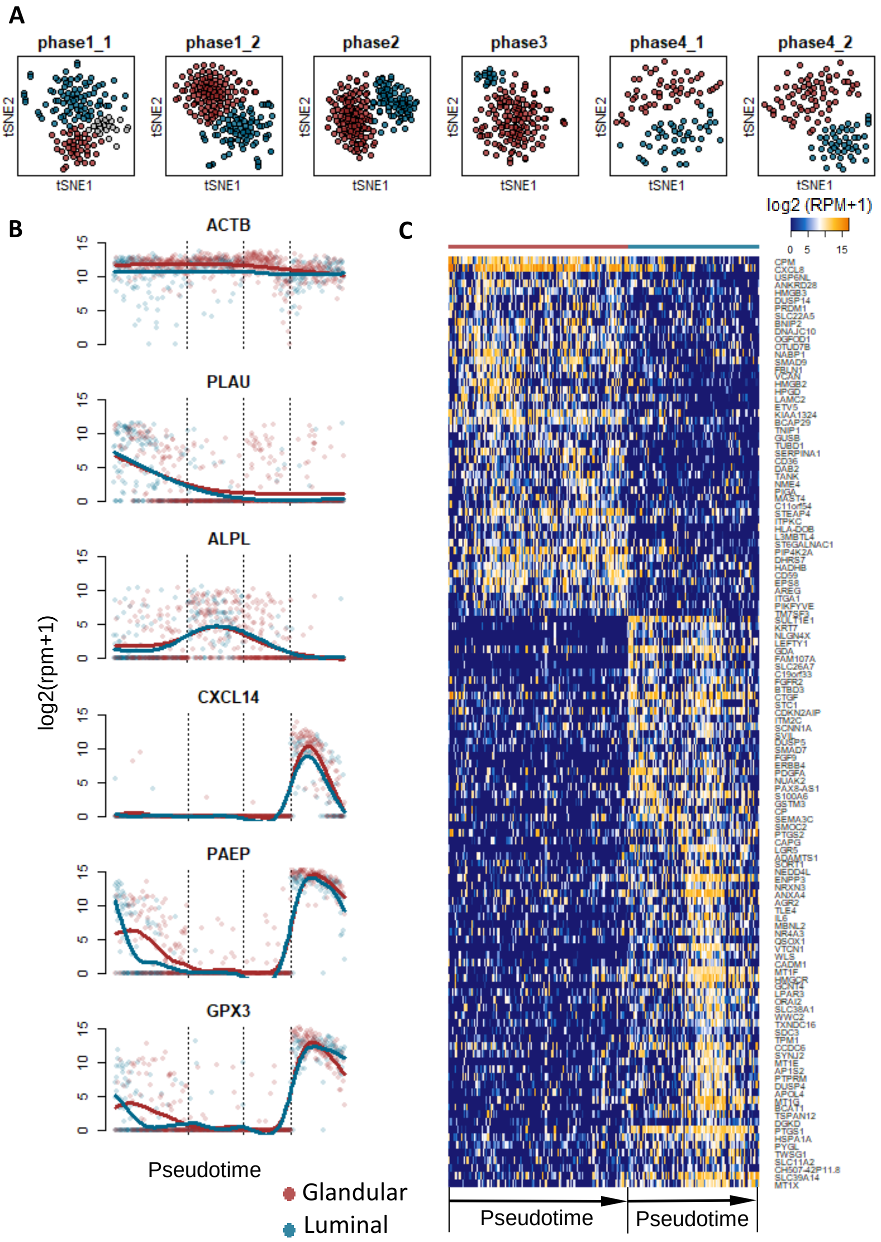


**Figure S10. Deviating subpopulations of unciliated epithelia and their transcriptomic signature across the human menstrual cycle**

(A) Dimension reduction (tSNE) on unciliated epithelia in major identified phases/sub-phases across the menstrual cycle

(B) Dynamics of phase-defining and housekeeping genes in unciliated epithelial subpopulations across the menstrual cycle (dashed lines: boundaries between the 4 phases)

(C) Dynamics of differentially expressed genes between the two sub-populations in phase 2

(Related to Figure 5)

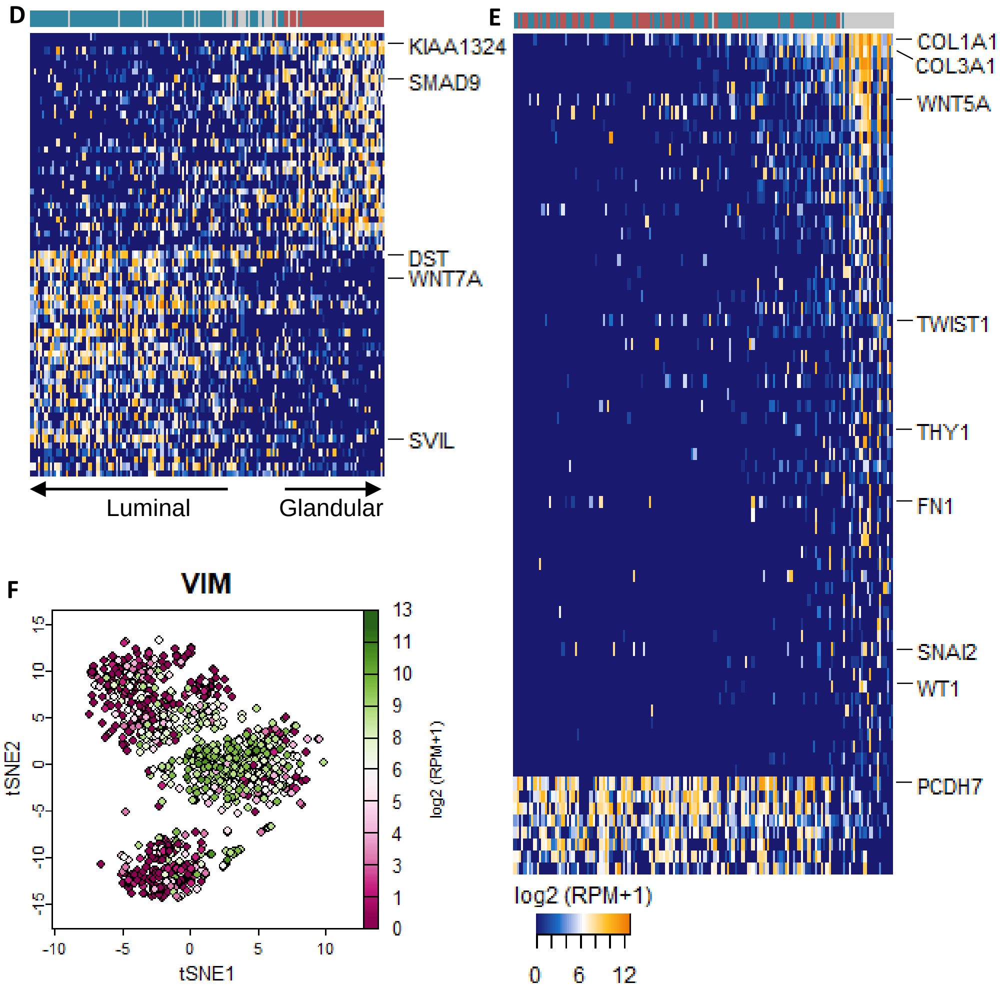


**Figure S10. (Continued)**

(D) Relationship between the ambiguous cell population with luminal and glandular cells in early phase 1. Shown are differentially expressed genes ($-\log_{10}$(p_adj of a Wilcoxon's rank sum test)>0.05, $\log_2$(FC)>2) between luminal and glandular epithelia in early phase 1. Cells (column) are ordered by the ratio of (average expression of genes upregulated in the luminal) and (average expression of genes upregulated in the glandular)

(E) Genes over-expressed and under-expressed in the ambiguous cell population relative to in luminal and glandular epithelia in early phase 1. Cells (column) are ordered by the ratio of (average expression of genes under-expressed) and (average expression of genes over-expressed)

(F) Expression of vimentin (VIM) in unciliated epithelia

(Related to Figure 5)

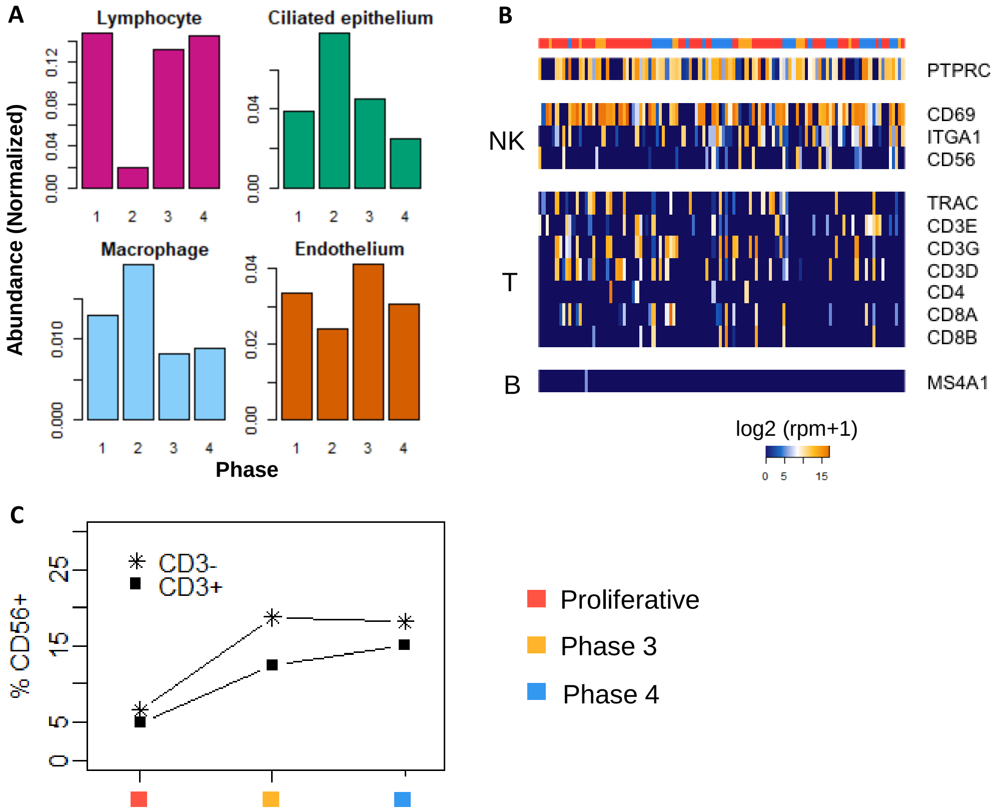


**Figure S11. Changes in other endometrial cell types across the human menstrual cycle**

(A) Normalized abundance of other endometrial cell types demonstrated phase-associated dynamics. Normalization was done against the total number of unciliated epithelia (for ciliated epithelium) or stromal fibroblasts (for lymphocyte, endothelium, macrophage) captured for each biopsy

(B) Expression of major lymphoid lineage markers. Cells (columns) were sorted based on percent NK receptors expressed (as in Figure 6A)

(C) Percent CD56+ cells in all "CD3+" and "CD3-" lymphocytes across major phases of the cycle

(Related to Figure 6)

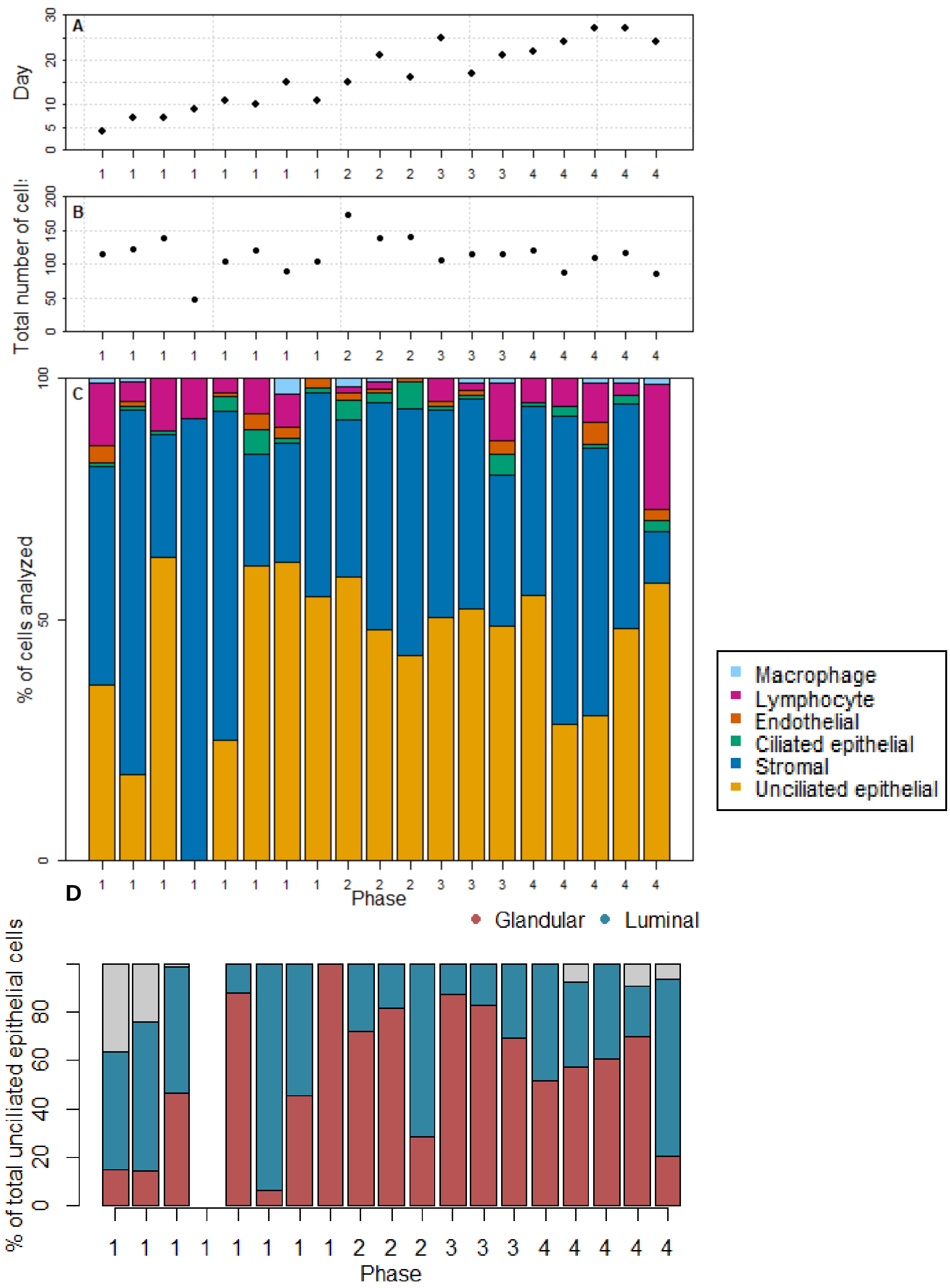


**Figure S12. Data summary**

For each woman:

(A) Relationship between the day of menstrual cycle and her assignment to one of the four major phases (Figure 3) based on unbiased single cell analysis

(B) Total number of single cells analyzed

(C) Distribution of the six cell types

(D) Distribution of glandular and luminal epithelia (Gray: cells in the ambiguous cell population in Figure 5A)

(Each dot (A, B) or each bar (C, D) represents a woman. From left to right, women were ordered, based on the median pseudotime of her stromal fibroblasts and unciliated epithelia)
